# Mechanisms of Resistance to CAR-T cell Immunotherapy: Insights from a Mathematical Model

**DOI:** 10.1101/2023.05.01.538974

**Authors:** Daniela S. Santurio, Emanuelle A. Paixão, Luciana R.C. Barros, Regina C. Almeida, Artur C. Fassoni

## Abstract

Chimeric Antigen Receptor (CAR)-T cell therapy long-term follow-up studies revealed non-durable remissions in a significant number of patients. Some of the mechanisms underlying these relapses include poor CAR T cell cytotoxicity or persistence, as well as antigen loss or lineage switching in tumor cells. In order to investigate how antigen-mediated resistance mechanisms affect therapy outcomes, we develop a mathematical model based on a set of integral-partial differential equations. Using a continuous variable to describe the level of antigen expression of tumor cells, we recapitulated important cellular mechanisms across patients with different therapeutic responses. Fitted with clinical data, the model successfully captured the dynamics of tumor and CAR-T cells for several hematological cancers. Furthermore, the role played by these mechanisms are explored with regard to different biological scenarios, such as pre-existing or acquired mutations, providing a deeper understanding of key factors underlying resistance to CAR-T cell immunotherapy.

**Statement of significance:** Our study introduces the first mathematical model to characterize the influence of a continuous level of antigen expression on the interplay between Chimeric Antigen Receptor (CAR)-T cells and cancer cells. We examine various cellular mechanisms across different hematologic cancers, taking into account both antigen-positive and antigen-negative relapses. Our findings shed light on the role of antigen density in CAR-T cell therapies and provide a valuable framework to investigate resistance with potential to improve patient’s outcomes.

## 1 Introduction

Chimeric antigen receptor (CAR)-T cell treatment is an immunotherapy in which T lymphocytes are genetically modified to interact with tumor cells upon antigen binding, triggering a powerful cytolytic response. In the past years, remarkable responses to the infusion of CAR-T cells in patients with relapsed and refractory hematological diseases led to the FDA’s approval of multiple products targeting cancer cells expressing CD19 and BCMA antigens [1, 2, 3, 4, 5].

Despite promising results, follow-up data show that maintaining long-term remission is a significant challenge in CAR-T cell therapy. While complete remission can be achieved in 70-90% of pediatric and adult ALL patients, long-term studies show that 30-60% of patients relapsed, the majority occurring one year after treatment [6, 7]. Thus, a deeper understanding of the factors associated with primary and acquired resistance to CAR-T therapy is necessary to decrease relapse rates and improve treatment efficacy [8].

CAR-T cells’ functionality is mediated by antigen recognition. Several biological mechanisms, such as expansion, cytotoxicity, and memory pool formation are influenced directly or indirectly by antigen receptor signaling. Thus, the antigen density presented by tumor cells is a major factor influencing CAR-T cell activity and leads to two patterns of therapy resistance: antigen-positive and antigen-negative relapses. Antigen-positive relapses are typically associated with tumor cells expressing high levels of antigen on their surface and are explained by limited persistence or low potency of CAR-T cells. In antigen-negative relapses, tumor cells no longer display antigen on their surface, resulting in tumors that evade CAR-mediated recognition and clearance, despite CAR T-cell persistence. Different mechanisms may underlie this type of relapse, including antigen loss due to genetic mutations [9], the existence of resistant clones, the pre-therapy heterogeneity in the antigen density expressed by tumor cells, resulting in the selection of resistant clones [10], lineage switch induced by immune pressure [11], transitory antigen loss due to CAR-T cell contact, in a process called trogocytosis [12] or due to non-mutational mechanisms, such as post-transcriptional factors and the generation of alternative isoforms [13, 14]. Although these mechanisms are known for leading to antigen-negative relapses, they are not fully elucidated and require further exploration.

Extensive research has been conducted in an attempt to improve the efficacy of CAR-T cells and overcome resistance mechanisms. Patients undergoing CAR-T immunotherapy already have intrinsic risk factors that cause defects in their T cells, which, when combined with multiple prior treatments at high drug doses, may result in a poor CAR product in terms of quality and quantity. Manufacturing CAR-T cells from healthy donors (‘off-the-shelf’ CAR-T) [15], creating a product rich in a memory-like phenotype [16] and, developing new CAR designs with combined co-stimulatory domains [17] or “armored CAR T-cells” that constitutively secrete active cytokines [18, 19] are all promising approaches. Further, sequential infusions, multi-antigen targeting, and combining CAR-T with other therapies (allo-HSCT, checkpoint inhibitors) are also new avenues to explore to enhance CAR-T effectiveness [20, 21]. Despite these efforts, the clinical difference in progression-free or overall survival is controversial and numerous questions regarding CAR-T cell resistance remain unanswered. Specially in the case of antigen loss, the underlying mechanisms vary, posing a new set of challenges for CAR-T cell treatment [7].

As more data gathered from clinical trials became available over the years, there has been a surge of interest in using mathematical models to describe different aspects of CAR-T therapy. These models are valuable tools for predicting different outcomes through *in silico* simulations and are among the most prominent methods used to study biological systems. Using clinical data, Hardiansyah and Ng [22] demonstrated connections between dose and disease burden by investigating the interplay of proinflammatory cytokines. Stein et al. [23] proposed the first mixed-effect model considering different CAR-T phenotypes by investigating the impact of co-medications for treating cytokine release syndrome after CAR-T cell expansion. In a preclinical model for hematological cancers, Barros et al. [24] modeled the dynamics of two CAR-T cell populations (effector and memory), highlighting the importance of the memory pool formation for the response to therapy. Subsequently, Paixão et al. [25] extended this model to the clinical scenario, resulting in a model that describe the patient-specific multiphasic dynamics of CAR-T cell kinetics through the differentiation of functional, memory, and exhausted phenotypes, as well as the dynamics of cancer cells. Singh et al. [26] used a multiscale PK/PD model to study the relationship between CAR-affinity, antigen abundance, tumor cell depletion, and CAR T-cell expansion in mouse models. Other mechanisms have been explored over the past years, for example, the interactions between CAR-T, cancer and healthy cells to predict efficacy and toxicity [27, 28], the efficacy of patient preconditioning lymphodepletion therapy [29], and the relation of CAR-T intracellular modulations and cellular response signaling [30]. There are also models for solid tumors [31, 32, 33, 34] and, up-to-date, Liu et al. proposed the only model that explored the interaction of activated/non-activated CAR-T and CD19+/CD19-tumor cells for patients with different therapy outcomes [35]. Reviews on modeling CAR-T cell therapy are found in [36, 37, 38].

Although the existing models are able to capture CAR-T dynamics and provide useful insights, none of them focused on how antigen density affects cellular mechanisms underlying heterogeneous responses. Here, we propose a mathematical model to investigate how antigen-mediated resistance affects treatment efficacy. We aim to capture different processes involved in the development of resistance to CAR-T cell immunotherapy, including transient antigen loss of tumor cells due to treatment pressure and permanent antigen loss due to mutations aiming to elucidate the specific contribution of antigen density on CAR-T cell dynamics under different biological scenarios.

## 2 Materials and Methods

### 2.1 Mathematical model

We propose a model for the interactions between CAR-T and tumor cells, considering two phenotypes for CAR-T cells, effector (*C*_*T*_) and memory (*C*_*M*_) cells, as well as two subpopulations for tumor cells, antigen-positive (*T*_*P*_) and antigen-negative (*T*_*N*_) cells. The schematic description of the model is shown in Figure 1a.

**Figure 1:**
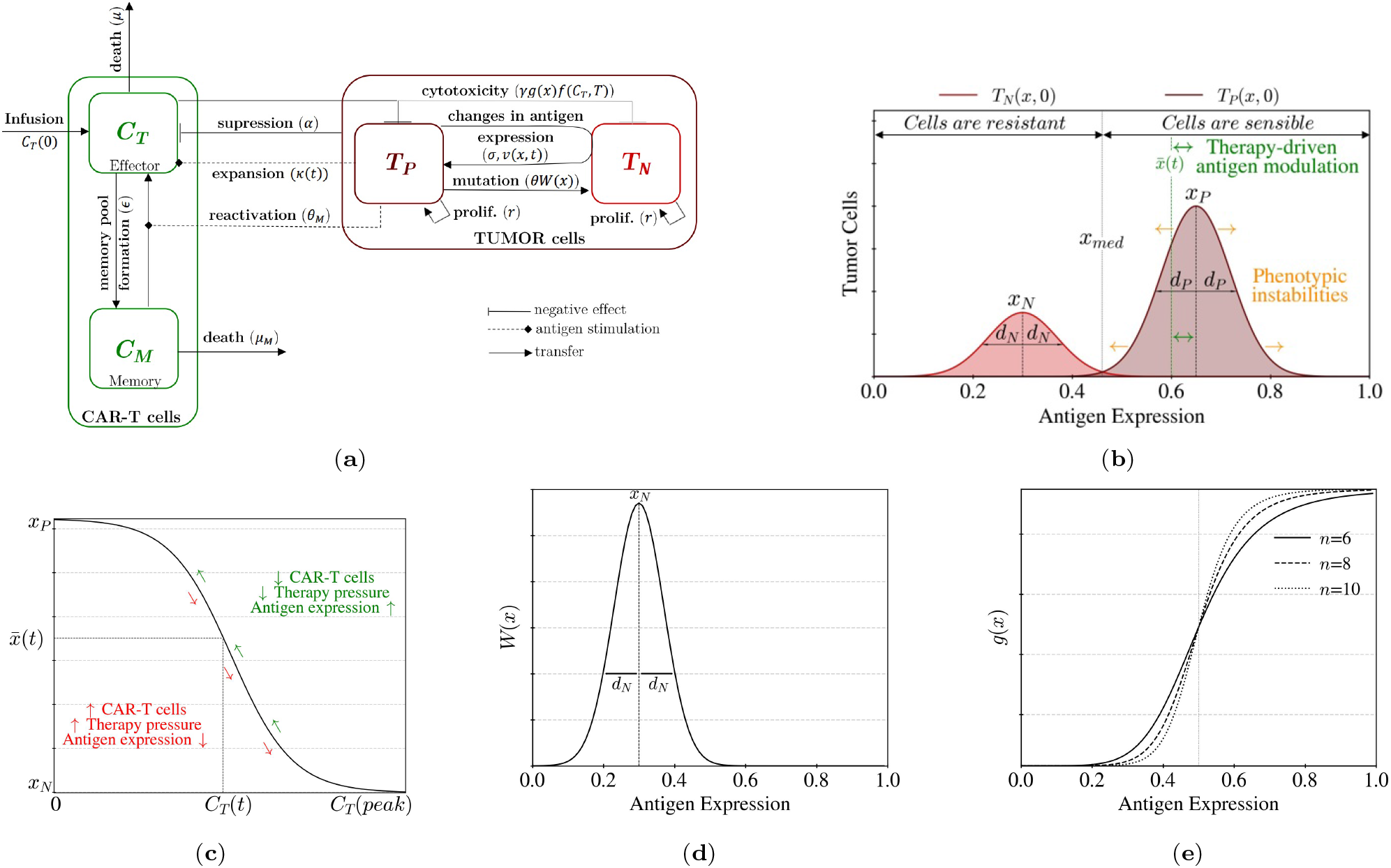
Modeling resistance mechanisms in CAR-T therapy. (a) Schematic description of the model structure. After infusion, effector CAR-T cells (*C*_*T*_) expand in response to antigen contact, differentiate into a memory phenotype, are suppressed by tumor cells, and undergo natural cell death. Memory cells (*C*_*M*_) have a longer persistence but may also die and differentiate back into effector CAR-T cells upon antigen re-exposure.Tumor cells are split into two constitutively different states: antigen-positive (Ag+, *T*_*P*_) and antigen-negative (Ag−, *T*_*N*_). All tumor cells proliferate but only Ag+ cells mutate into Ag− cells and experience transient changes in antigen expression. Since CAR-T cytotoxicity requires antigen-binding, tumor cells are killed with different rates, which depend on antigen-expression levels. (b) The initial distributions of tumor cells along the antigen space are given by Gaussian distributions with means *x*_*N*_ and *x*_*P*_, representing the homeostatic values of each population antigen-density, and *d*_*N*_ and *d*_*P*_ are the intrinsic variabilities around this level (standard deviations). The threshold of antigen detection, *x*_*med*_, defines two regions of therapy responses: within the region 0 ≤ *x* < *x*_*med*_, tumor cells are considered resistant, while within the region *x*_*med*_ ≤ *x* ≤ 1, they are considered sensible to CAR-T cell therapy. (c) Treatment pressure induces changes in antigen expression. This is modeled by the therapy-driven level of antigen-expression 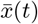 and its dependence on the CAR-T cell load. During CAR-T cells expansion 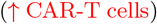, antigen-positive cells lose antigen and escape immune response 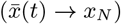. As the number of CAR-T cells decreases 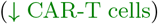, these cells can restore their antigen expression to its homeostatic level 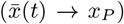. (d) The mutation kernel *W* (*x*) is defined as a Gaussian distribution with mean *x*_*N*_ and standard deviation *d*_*N*_ and denotes the probability that an antigen-positive tumor cell will mutate into a antigen-negative tumor cell with antigen expression *x*. (e) CAR-T cytotoxic function. The killing of tumor cells by CAR-T cells is regulated by antigen receptor signaling and, since tumor cells display heterogeneous antigen expression, the function *g*(*x*) represents the CAR-T cell cytotoxicity as an antigen-driven mechanism.

First, we describe the dynamics of the tumor cells. Experimental analysis by flow cytometry often shows antigen density distributions of tumor cells as a two-component log-normal function [39]. To address this heterogeneity in antigen expression, we follow the modeling approach of [40] and use a continuous variable *x* ∈ [0, 1] to represent antigen density, which is a normalization of the original antigen expression measures (see Section SM-4.1). Then, the two distinct genotypes, *T*_*P*_ (*x, t*) and *T*_*N*_ (*x, t*), represent the antigen-positive (Ag+) and antigen-negative (Ag−) tumor cells with antigen expression level *x* at time *t*.

While antigen-positive and antigen-negative tumor cells are constitutively distinct, we assume that Ag+ cells may experience temporary loss of antigen upon CAR-T cell contact [12] or due to post-transcriptional mechanisms [13, 14]. Thus, we define a threshold of the antigen expression *x*_*med*_ that splits the phenotypic space into two regions: cells with expression below this threshold of detection (0 *≤ x < x*_*med*_) are considered resistant to CAR-T cell therapy, while cells with expression above it (*x*_*med*_ *≤ x ≤* 1) are considered sensible (Figure 1b). Therefore, Ag+ cells that are initially sensible to treatment may become temporarily resistant, due to the phenotypic changes that reduce their antigen expression. Thus, the number of tumor cells that are sensible to CAR-T cell therapy is defined as

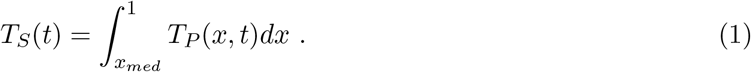

Following previous approaches [41, 40], we describe the dynamics of tumor cells with the following system of integral-partial differential equations:

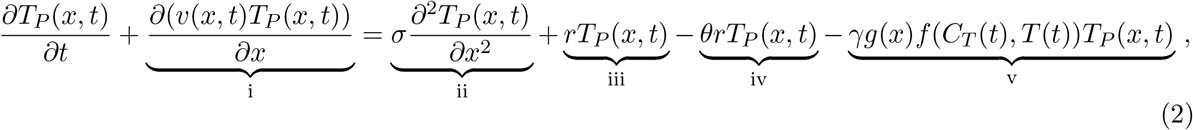

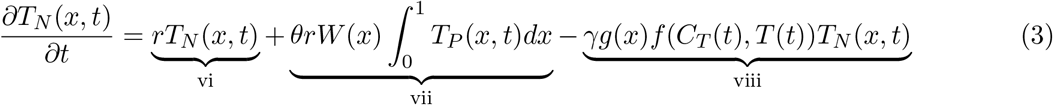

where the total tumor cell population is given by

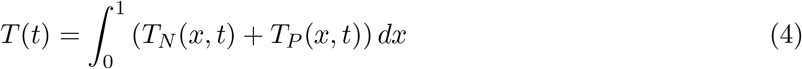

and the initial distributions are given by

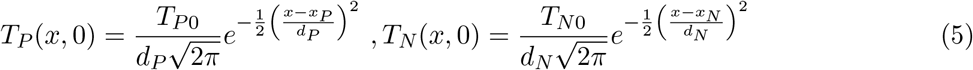

where *T*_*P*0_ and *T*_*N*0_ are the initial number of antigen-positive and antigen-negative tumor cells, respectively. As displayed in Figure 1b and specified in Equation (5), the distributions are defined in terms of their homeostatic mean antigen expression level (*x*_*N*_ and *x*_*P*_), as well as their corresponding standard deviations (*d*_*N*_ and *d*_*P*_). Details regarding the parameters of the initial distributions can be found in Section SM-4.1.

Equations (2) and (3) assume that tumor cells may experience two types of phenomena: conservative interactions, that encompass all processes involving changes in antigen expression while preserving the overall tumor cell population (terms (i) and (ii)); and non-conservative interactions, which include all processes that do not preserve the overall tumor cell population (terms (iii)-(viii)).

We assume that the advection of antigen-positive tumor cells along the antigen space (term (i)) is related to a therapy-driven level of antigen expression 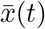 that encompasses a reversible antigen loss as a consequence of post-transcriptional mechanisms. Specifically, we assume that the antigen expression of Ag+ tumor cells under therapeutic pressure changes according to the expression:

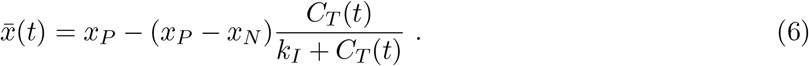

This formulation implies that, prior to the therapy, when there are no CAR-T cells in the system (*C*_*T*_ = 0), the therapy-driven level of antigen expression equals the homeostatic mean expression level of Ag+ cells, *x*_*P*_, while once therapy is started, CAR-T cells expand, and 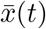 decreases progressively until reaching its minimum value, close to the homeostatic expression level of Ag− cells, *x*_*N*_ (Figure 1c). Finally, when CAR-T cells enter the contraction phase, this process reverses, and 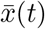 gradually increases until returning back to *x*_*P*_. The half-saturation constant *k*_*I*_ regulates this phenotypic transition. Higher values of *k*_*I*_ are associated with a slower rate of change in antigen expression, implying that tumor cells are less exposed to post-transcriptional events.

Therefore, the advective term (i) in Equation (2) describing the phenotypic drift along the antigen space is always directed towards the therapy-driven level of antigen expression 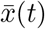 and is formally defined as

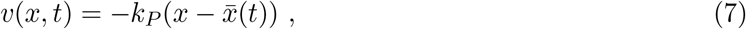

in which the rate *k*_*P*_ modulates the intensity of the advective flux and is named as the phenotypic transition coefficient of antigen expression. Thus, when therapy decreases the therapy-driven level 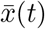, the distribution of Ag+ cells, *T*_*P*_ (*x, t*), is driven towards the region of low antigen-expression, which makes them temporarily resistant to therapy. Eventually, without the pressure of therapy, 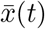 increases until reaching the homeostatic level *x*_*P*_ and the drift velocity enables Ag+ cells to restore their antigen expression.

The diffusive term (ii) in Equation (2) describes phenotypic instabilities in antigen expression. Following [42], we employ a diffusion operator to simulate small phenotypic changes induced by epigenetic factors. For a more detailed explanation about the conservative parts of equations (2) and (3) and its steady-state solution, see Section SM-4.2.

Regarding the non-conservative mechanisms in Equations (2) and (3), it is assumed that: (iii) and (vi) tumor cells proliferate at a rate *r*; (iv) and (vii) antigen-positive tumor cells mutate to an antigen-negative genotype during clonal division with rate *θ*; (v) and (vii) tumor cells are killed by CAR-T cells at rate *γg*(*x*)*f* (*C*_*T*_, *T*). It is important to note that proliferation occurs without changing the antigen expression but mutations may drive antigen loss in a one-way process. Specifically, a key assumption is that mutation-mediated genetic jumps on antigen density are allowed from Ag+ to Ag− tumor cells but not backward. As in [40], the parameter *θ ∈* [0, 1] indicates the fraction of tumor cells that experience mutation, so that (1 – *θ*) denotes the fraction of cells undergoing true division. The mutation kernel *W* (*x*) is defined as a Gaussian distribution with mean *x*_*N*_ and standard deviation *d*_*N*_ (Figure 1d),

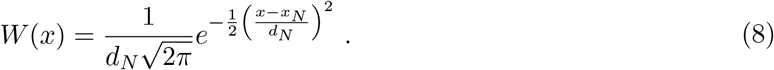

The killing of tumor cells by CAR-T cell (terms (v) and (vii)) occurs at a rate *γ*, and depends on the antigen density (as described by the function *g*(*x*)) and on the ratio between *C*_*T*_ (*t*) and *T* (*t*) (expressed by the function *f* (*C*_*T*_, *T*)). The antigen-dependent cytotoxicity function *g*(*x*) and the lysis function *f* (*C*_*T*_, *T*) are respectively given by:

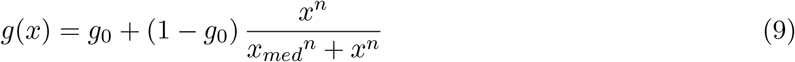

and

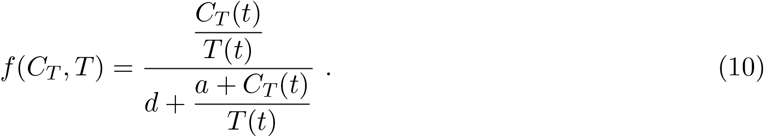

Here, *f* (*C*_*T*_, *T*) is a modified Hill function inspired by [43, 25]. For the cytotoxicity function, we propose a smooth step function *g*(*x*) *∈* [*g*_0_, 1] that modulates CAR-T cell cytotoxicity as an antigen-driven mechanism. As displayed in Figure 1e, (1 *− g*_0_) defines the height of the step, and *n* controls the function sharpness. When 0 *≤ x < x*_*med*_ (low antigen expression), CAR-T cells eliminate tumor cells through antigen-independent mechanisms at a small rate of *γg*_0_, while for *x*_*med*_ *≤ x ≤* 1 (high antigen expression) the maximum cytotoxicity rate of CAR-T cells, *γ*, is reached. A positive minimum fraction *g*_0_ *>* 0 is taken by considering killing mechanisms like those due to the bystander effect, CAR-T endogenous TCR, and other pro-inflammatory molecules [44].

Now, we describe the CAR-T cell population. Divided into two phenotypically distinct subpopulations, with *C*_*T*_ (*t*) representing the cytotoxic effector and *C*_*M*_ (*t*) for the memory CAR-T cells, their dynamics are based on our previous model [24] and is described by:

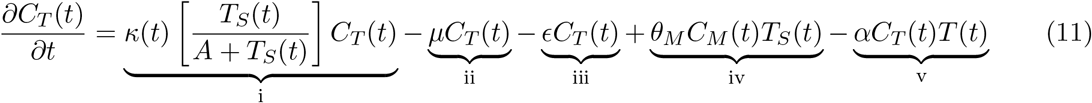

and

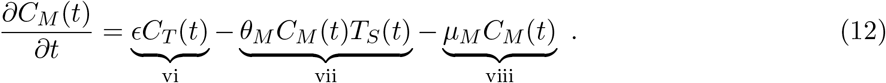

Equation (11) assumes that effector CAR-T cells (i) undergo clonal expansion with a time-dependent rate *κ*(*t*) after binding with sensible tumor cells (*T*_*S*_(*t*)); (ii) die naturally at a rate *μ*; (iii) differentiate to a memory phenotype at a rate *ϵ*; (iv) receive re-activated memory cells due to antigen re-exposure at a rate *θ*_*M*_ *T*_*S*_(*t*); and (v) are inhibited by tumor-induced immunosuppressive effects at a rate *αT* (*t*).

Based on our previous work [25], the expansion function *κ*(*t*) displays a high expansion level that may be sustained for a certain period, forming a plateau, before finally decreasing to a baseline level. This is formally described by

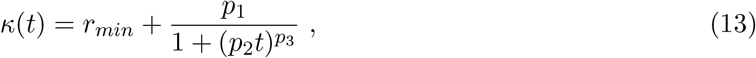

where *r*_*min*_ is the basal CAR-T cell expansion rate, *p*_1_ defines the initial expansion rate, while *p*_2_ and *p*_3_ modulate the duration and the decay of the expansion rate, respectively. These are patient-specific parameters and describe heterogeneous patterns of CAR-T cell proliferation.

Equation (12) describes the dynamics of memory CAR-T cells and assumes that (vi) memory CAR-T cells are formed from the differentiation of effector CAR-T cells at a rate *ϵ*, (vii) upon antigen contact, memory cells undergo a fast increase in metabolic activity and immediately return to the effector CAR-T cell phenotype with rate *θ*_*M*_ [45], and (viii) die naturally at a rate *μ*_*M*_.

Descriptions of all model parameters and corresponding units are given in Table 1, while details regarding the model’s biological assumptions are in Section SM-1.

**Table 1:** Model parameters and functions with their units and biological meaning.

| Parameter | Unit | Biological meaning |
| --- | --- | --- |
| $r$ | $\text{day}^{-1}$ | Growth rate of tumor cells |
| $\theta$ | — | Mutation rate from antigen-positive tumor cells to antigen-negative tumor cells |
| $A, a$ | cell | Half-saturation constants of CAR-T cell expansion and lysis function |
| $d$ | — | Half-saturation constant of the cytotoxic effect on tumor cells |
| $k_P$ | $\text{day}^{-1}$ | Phenotypic transition coefficient of antigen expression |
| $\sigma$ | $\text{day}^{-1}$ | Diffusion operator that represents the phenotypic changes induced by epigenetic factors |
| $x_P, x_N$ | — | Homeostatic mean antigen expressions of antigen-positive and antigen-negative tumor cells |
| $d_P, d_N$ | — | Intrinsic variability of initial antigen-positive and antigen-negative tumor cells |
| $x_{med}$ | — | Threshold of antigen detection |
| $n$ | — | Hill coefficient of the antigen-mediated cytotoxicity function $g(x)$ |
| $k_I$ | cell | Saturation constant that modulates the therapy-driven phenotypic transition of tumor cells |
| $\gamma$ | $\text{day}^{-1}$ | Maximum cytotoxic rate of effector CAR-T cell |
| $g_0$ | — | Minimum fraction of CAR-T cytotoxicity |
| $\alpha$ | $(\text{cell} \cdot \text{day})^{-1}$ | CAR-T cell inactivation/inhibition coefficient by tumor cells |
| $\mu$ | $\text{day}^{-1}$ | Death rate of effector CAR-T cells |
| $r_{min}$ | $\text{day}^{-1}$ | Minimum expansion rate by effector CAR-T cells |
| $p_1$ | $\text{day}^{-1}$ | Initial expansion rate by effector CAR-T cells |
| $p_2$ | $\text{day}^{-1}$ | Rate that regulates the duration of maximum expansion period of effector CAR-T cells |
| $p_3$ | — | Expansion coefficient that regulates the decay of maximum expansion period of effector CAR-T cells |
| $\epsilon$ | $\text{day}^{-1}$ | Conversion rate of effector CAR-T cells into memory CAR-T cells |
| $\theta_M$ | $(\text{cell} \cdot \text{day})^{-1}$ | CAR-T cell conversion coefficient from memory to effector phenotype due to contact with sensible tumor cells |
| $\mu_M$ | $\text{day}^{-1}$ | Death rate of memory CAR-T cells |
| Function | Unit | Biological meaning |
| $\kappa(t)$ | $\text{day}^{-1}$ | CAR-T cell expansion function |
| $\bar{x}(t)$ | — | Therapy-driven level of antigen expression |
| $v(x, t)$ | $\text{day}^{-1}$ | Drift velocity |
| $W(x)$ | — | Mutation kernel |
| $g(x)$ | — | Antigen-dependent cytotoxicity function |
| $f(C_T, T)$ | — | Lysis function |

### 2.2 Patient data, model fitting and numerics

We gathered CAR-T cell counts from 18 patients [46, 47, 10, 48, 49] with various cancer types and fitted the model parameters to display the data-informed therapy responses. Patients were grouped according to their outcome, i.e., Complete Response (CR), Antigen-Positive Relapse (*Relapse*+), and Antigen-Negative Relapse (*Relapse*−). Detailed information on clinical data, patient-specific parameter estimation, and the numerical simulation of the model are given in Sections SM-2 to SM-6.

### 2.3 Possible scenarios leading to resistance

To investigate the key processes involved in the emergence of relapse, we assumed two scenarios:

- SC_1_ - Pre-Existing Antigen-Negative Cells: There is one Ag− cell within the initial tumor burden (*T*_*N*0_ = 1 cell) and no genetic mutations (*θ* = 0) occur during therapy.
- SC_2_ - Random Mutations After CAR-T Cell Therapy: Initially, there are no Ag− cells (*T*_*N*0_ = 0 cell) and the resistant population arises from genetic mutations (*θ* ≠ 0). To investigate the mutation rate (*θ*) variability between the cohorts, the minimum fraction of CAR-T cytotoxicity was fixed at *g*_0_ = 0.015.

## 3 Results

We model the interactions among antigen-positive (*T*_*P*_) and antigen-negative (*T*_*N*_) tumor cells and effector (*C*_*T*_) and memory (*C*_*M*_) CAR-T cells. Each tumor subpopulation exhibits phenotypic heterogeneity with respect to antigen expression *x ∈* [0, 1], which allows considering different resistance mechanisms such as trogocytosis, post-transcriptional antigen loss, and mutations. Using the parameter values displayed in Table SM-4, we present our main findings in this section.

### 3.1 Model assessment for patients with different therapy outcomes

We first evaluated how the proposed model describes the dynamics in patients with the distinct responses *Relapse*−, *Relapse*+, and *CR*. We considered that all 18 patients received a single infusion of CAR-T cells on day zero and assumed a baseline tumor burden of *T*_*P*0_ = 10^7^ cells. Specifically for patient G02, the initial distribution of antigen expression was available, consisting of 7.7% antigen-negative cells and 92.3% antigen-positive cells (see Section SM-2). Since the proposed scenarios produce similar outcomes, Figure 2 displays the simulations for SC_1_ while those for SC_2_ are displayed in Figure SM-4.

**Figure 2:**
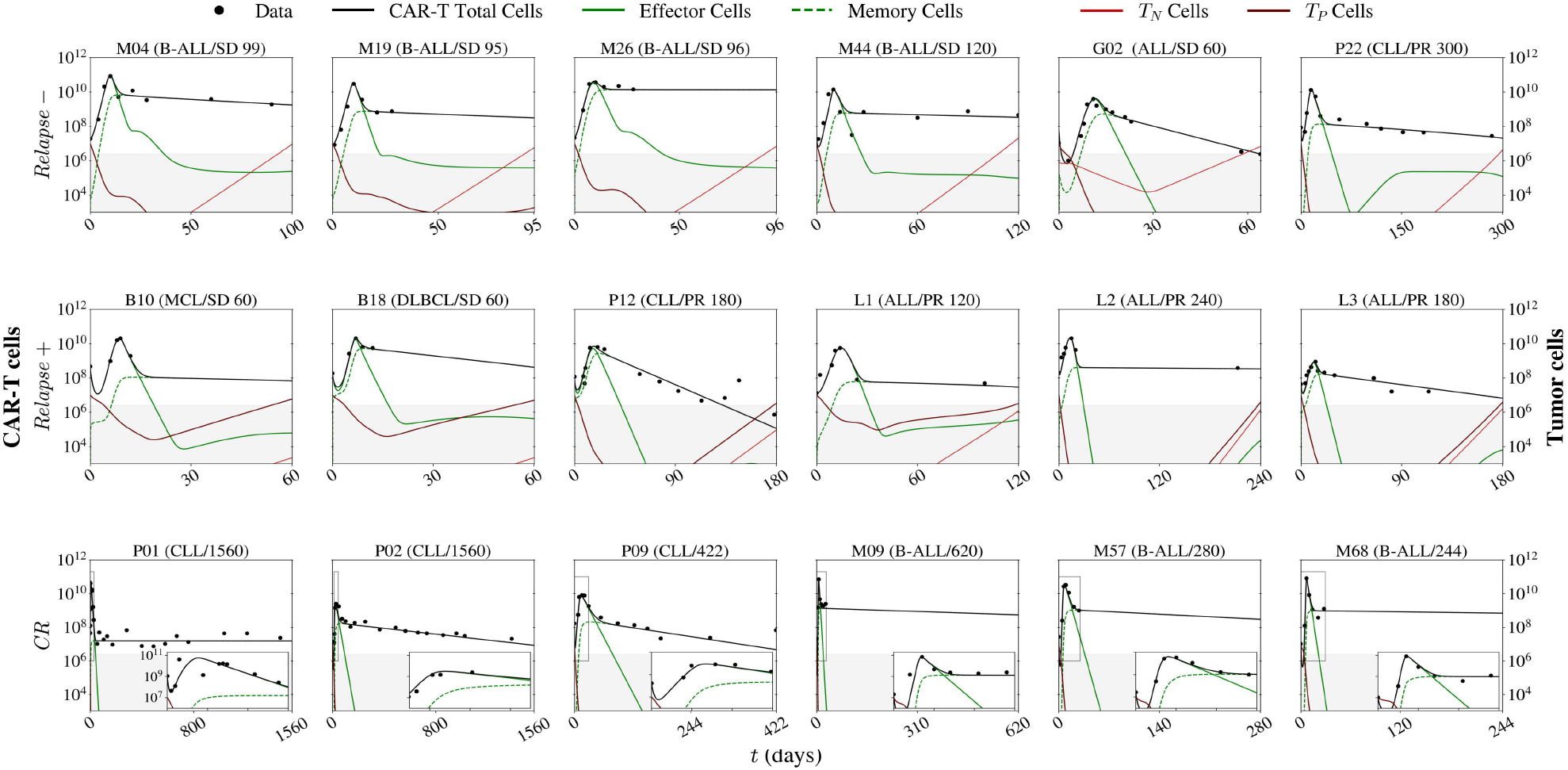
Model fits of tumor-CAR-T cell interactions for 18 patients divided into three cohorts: *Relapse*− (top panel), *Relapse*+ (middle panel), and *CR* (bottom panel). The total CAR-T cell population includes effector (*C*_*T*_) and memory (*C*_*M*_) phenotypes, while the total tumor population encompasses antigen-negative (*T*_*N*_) and antigen-positive (*T*_*P*_) cells. The qPCR detection threshold of 2.5 × 10^6^ cells was represented by the gray area. The zoomed-in analysis of patient P02 was made from day 15 to day 45, while the remaining CR patients were analyzed from day 0 to day 30. B-ALL, B-cell acute lymphoblastic leukemia; ALL, acute lymphoblastic leukemia; CLL, chronic lymphoblastic leukemia; MCL, mantle cell lymphoma; DLBCL, diffuse large B cell lymphoma. These simulations consider SC_1_ (pre-existing Ag− cells, no mutations); corresponding fits for SC_2_ are presented in SM-4

The model has successfully captured the experimentally observed CAR-T kinetics, in which CAR T-cells rapidly expand upon injection and reach their peak within the first 2 weeks. Then, the effector CAR-T cells rapidly decrease during the contraction phase until the total population is predominantly formed by the long-lasting memory cells, defining the persistence phase [25]. Most of the patients in the *Relapse*− cohort have a large memory pool, a marked persistence phase, and it is possible to observe a significant amount of CAR-T effector cells even if below the detection threshold (Figure 2, top panel). In the *Relapse*+ cohort, patients B10, L1, and L3, present a smaller pool of memory cells while patients P12 and L2 present significant depletion of effector cells (Figure 2, middle panel). Patients with *CR* display a large expansion of CAR-T cells followed by a long-lasting persistence phase (Figure 2, bottom panel). Furthermore, the model also captured sustained remissions in the *CR* cohort, early relapses (B10, B18, and L1), and longer remissions followed by late relapses (P12, L2, and L3). Finally, despite the pre-existence of negative tumor cells in patient G02 and a lineage switch in patients M26 and P22, the model fully reproduced their dynamics.

### 3.2 The interplay between tumor relapse and antigen expression

We now investigate the dynamics of tumor cells in response to antigen modulation. To do so, two representative patients, P22 and L2, were selected based on their treatment responses. For the *Relapse*− patient (P22), CAR-T cells eliminated all Ag+ cells even when antigen expression decreases (Figure 3a, *t* = 300). However, since a resistant cell arises, whether it was already present prior to treatment (SC_1_) or acquired through a mutation (SC_2_), the tumor manages to escape (Figure 3a, *t* = 130). On the other hand, for the *Relapse*+ patient (L2), despite a decrease in antigen expression during the first initial weeks, CAR-T cells are not able to eliminate all Ag+ cells.

**Figure 3:**
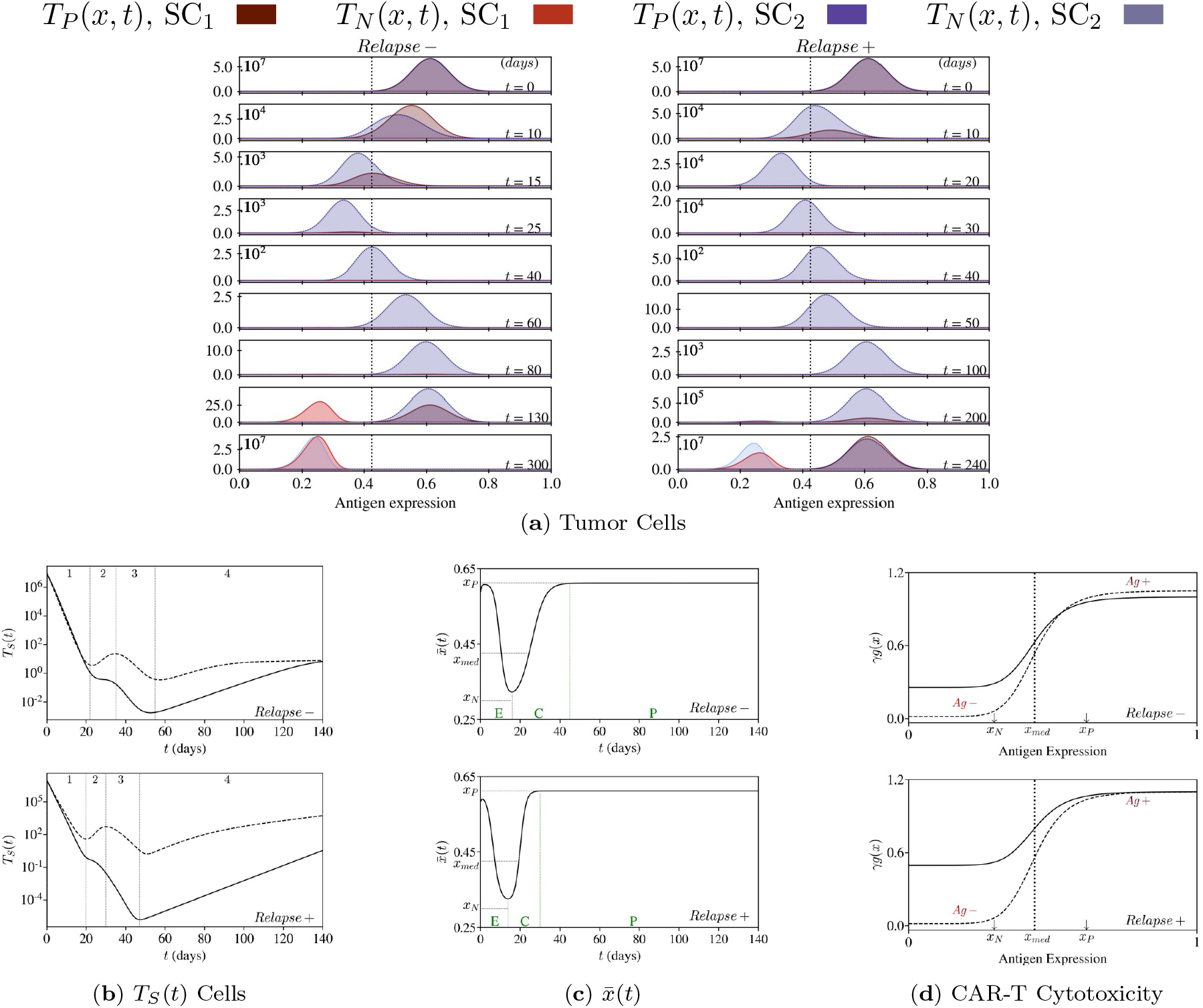
Tumor dynamics in response to antigen-modulation for the representative patients P22 (*Relapse*− panels) and L2 (*Relapse*+ panels) in scenarios 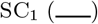 and 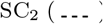. (a) Tumor dynamics along the antigen space. During the treatment, Ag+ tumor cells are killed by CAR-T cells and change their antigen expression. Mechanisms like genetic mutations, the pre-existence of antigen-negative clones, or CAR-T impaired cytotoxicity may lead to tumor progression. (b) Time evolution of tumor sensible cells (*T*_*S*_) is defined in four stages: 1) initial decay due to CAR-T cell expansion; 2) slower decay (SC_1_) and re-growth (SC_2_) due to antigen loss; 3) secondary decay during antigen restoration; and 4) fast (SC_1_) and slow (SC_2_) re-growth due to CAR-T persistence. (c) Time evolution of the therapy-driven level of antigen expression 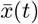. Periods of the duration of the expansion, contraction, and persistence phases of CAR-T cells are indicated by the letters E, C, and P, respectively. During the expansion phase (E) of CAR-T cells, driven by the therapy pressure, antigen-positive tumor cells lose antigen and evade the immune attack. As the number of CAR-T cells decreases in the contraction phase (C), antigen-positive tumor cells restore their homeostatic level of antigen expression. (d) CAR-T cell cytotoxicity depends on the antigen density. Tumor cells with antigen downregulation (*Ag−*) are less likely to be killed by antigen-independent mechanisms, while tumor cells displaying high amounts of antigen (*Ag*+) are more susceptible to being killed by CAR-T cells.

At the beginning of therapy, during CAR-T cell expansion, antigen-positive tumor cells are sensible (*T*_*S*_) and undergo depletion (Figure 3b, stage 1). As CAR-T cells reach their peak, antigen-positive tumor cells reach their lowest antigen expression level, with a value of 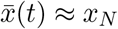. At this stage, these tumor cells are exhibiting a resistant phenotype (Figure 3c). Consequently, CAR-T cytotoxicity is severely impaired (Figure 3d, *Ag*−). Despite the presence of CAR-T cells in the system, antigen down-regulation opens a window for tumor re-growth, and if this condition persists, an early escape may occur (Figure 3b, stage 2). This dynamic is more pronounced for SC_2_ due to the reduced cytotoxicity of CAR-T cells towards tumor cells with a low antigen expression of 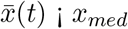. Specifically, patient L2 experiences faster changes in antigen expression (Figure 3c). Furthermore, to control the selection and growth of resistant clones, significantly higher levels of antigen-independent mechanisms (*g*_0_) are required, implying that the hypothesis of pre-existing resistant cells prior to treatment is less likely. Nevertheless, antigen-positive tumor cells (*T*_*P*_) are not completely eliminated, indicating poor CAR-T cell persistence or insufficient cytotoxicity towards tumor sensible cells (Figure 3d, *Ag*+). Lastly, while CAR-T cells are in the persistence phase, there are no further changes in antigen expression (Figure 3c), leading to two possible outcomes: (a) if a substantial number of sensible cells remain, the effector CAR-T cells can be re-stimulated. If not exhausted, they can control the emergence of a resistant population while killing the remaining antigen-positive cells; (b) if the residual number of sensible cells is limited, the effector CAR-T cells are less stimulated and may be insufficient to avoid tumor escape (Figure 3b, stage 4).

### 3.3 Mechanisms underlying resistance to CAR-T therapy

To investigate the interplay between the resistance mechanisms and CAR-T cell immunotherapy, we next analyze the influence of different tumor parameters on the system dynamics in both SC_1_ and SC_2_. To do so, we selected patients P22 and P12, who respectively experienced negative and positive relapses, and examined the influence of parameters *k*_*P*_, *k*_*I*_, and *d*_*P*_. The first parameter modulates the intensity of drift-velocity and phenotypic transition along the antigen space (see Equation (7)), while the second modulates the sensibility of therapy-driven antigen loss, in Equation (6). The third parameter represents the intratumoral heterogeneity of antigen expression within the antigen-positive population. Our simulations for both patients are presented in Figure 4.

**Figure 4:**
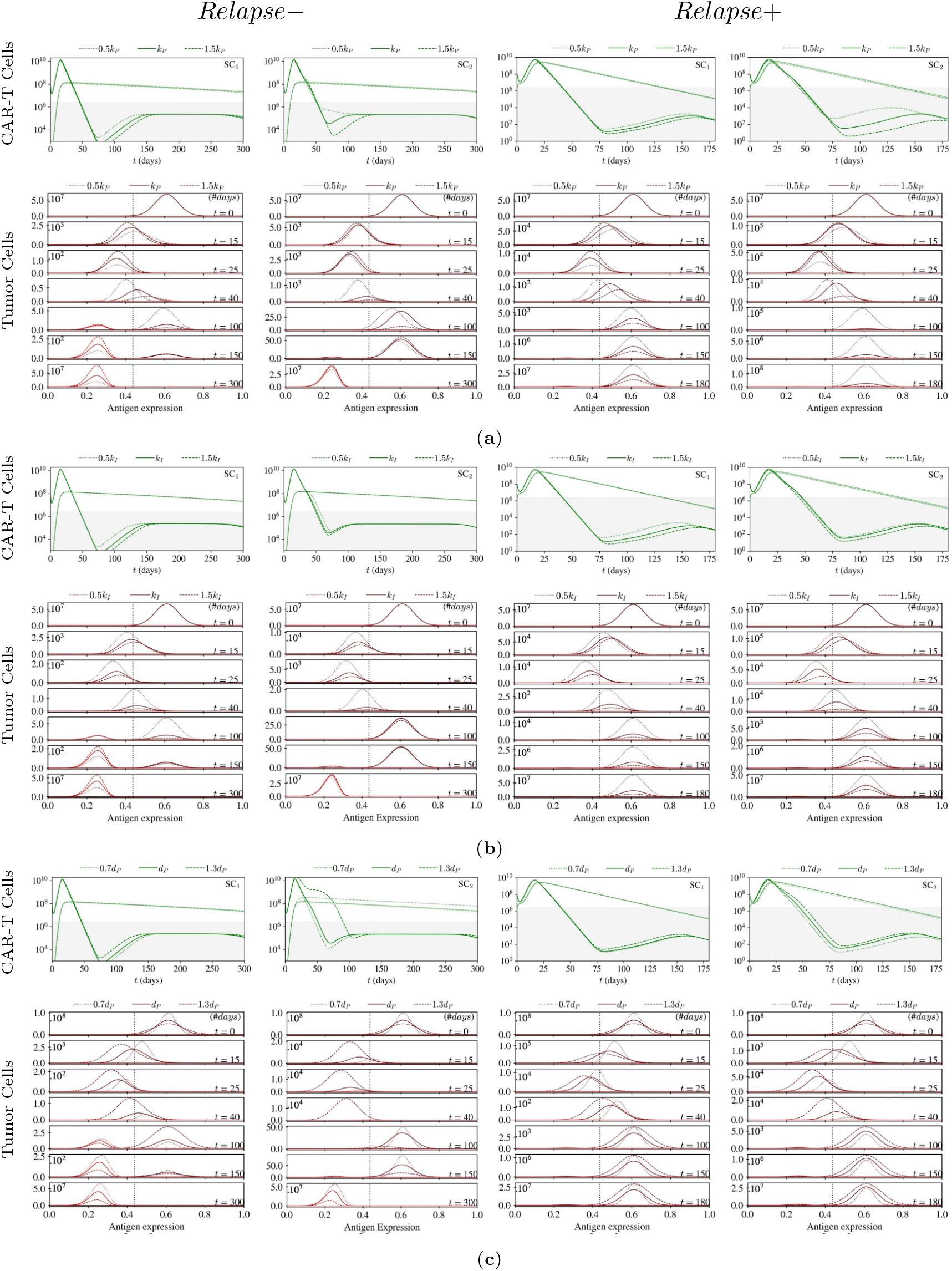
Model simulations for different resistance mechanisms for patient P22 (first and second columns) and patient P12 (third and fourth columns). After fitting the model, parameters related to (a) the phenotypic transition (*k*_*P*_), (b) the therapy-driven antigen loss (*k*_*I*_), and (c) the intratumoral heterogeneity of antigen-positive tumor cells (*d*_*P*_) were varied to assess changes in the dynamics of effector 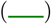 and memory 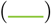 CAR-T cells, as well as changes in antigen-positive 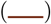 and antigen-negative 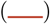 tumor cells. The corresponding scenarios are indicated in the CAR-T cell figures.

#### 3.3.1 Phenotypic transition and epigenetic instabilities of antigen expression (*k*_*P*_)

We first varied parameter *k*_*P*_, as displayed in Figure 4a. Our simulations indicate that changes in the coefficient of phenotypic transition *k*_*P*_ had minimal effect on CAR-T cell expansion and memory formation, but impacted effector cell depletion and re-expansion time span. As *k*_*P*_ decreases, conservative effects weaken and the drift velocity slows down. This results in tumor cells with a more heterogeneous phenotype (flattened aspect of *T*_*P*_ (*x, t*) distribution) and indicates that it takes longer for antigen-positive cells to transition back to a phenotypic region where they are sensible to the therapy.

A reduced phenotypic transition coefficient (0.5*k*_*P*_) for patient P22 resulted in a larger population of sensible cells in SC_1_, which enabled a greater stimulation of effector CAR-T cells, leading to better control of *T*_*N*_ cells and delayed relapse. In SC_2_, CAR-T effector cells suffer less depletion and had no effect on relapse time. For patient P12, the limited number of effector CAR-T cells or impaired cytotoxicity allowed sensible cells to evade CAR-T activity, especially in SC_2_. In summary, our findings suggest that a smaller value of phenotypic transition coefficient *k*_*P*_ delays the antigen-negative relapse and accelerates the antigen-positive relapse.

#### 3.3.2 Therapy-driven antigen loss (*k*_*I*_)

Figure 4b shows that therapy-induced changes in antigen expression (*k*_*I*_) had minimal effects on CAR-T cell dynamics, but impacted patient outcomes. In particular, Ag+ cells less susceptible to antigen down-regulation via post-transcriptional events (1.5*k*_*I*_) facilitated CAR-T cell binding due to a slower antigen loss, resulting in a faster antigen-negative relapse for P22 in SC_1_. In SC_2_ the variations evaluated for *k*_*I*_ had little effect on cell populations at the end of the simulation. A higher value of *k*_*I*_ slightly delayed relapse for P12 but didn’t affect initial tumor cell depletion.

Conversely, faster changes in antigen expression (0.5*k*_*I*_) allowed more Ag+ cells to become temporarily resistant and quickly transition back to a sensible phenotype, maintaining a larger population of these *T*_*S*_ cells. This led to a second expansion of effector CAR-T cells and delayed antigen-negative relapse for P22. However, for P12, it opened an escape window, resulting in a faster antigen-positive relapse. In summary, a smaller *k*_*I*_ value leads to faster therapy-induced changes in antigen expression, accelerating the antigen-positive relapse and delaying the antigen-negative relapse.

#### 3.3.3 Intratumoral heterogeneity of antigen-positive tumor cells (*d*_*P*_)

To assess the role of the antigen intratumoral heterogeneity, we hypothesized that antigen-positive tumor distribution would have the same homeostatic mean (*x*_*P*_) but different intrinsic variability of initial antigen expression (*d*_*P*_). Figure 4c shows that a more heterogeneous population of Ag+ cells (1.3*d*_*p*_) is advantageous for controlling the emergence of resistant clones but disadvantageous for controlling the proliferation of antigen-positive tumor cells. Indeed, a larger number of Ag+ cells undergo the phenotypic transition, which leads to a greater number of effector CAR-T cells in the system and delays the escape of tumor cells in patient P22. However, for patient P12, the stronger depletion of effector CAR-T cells promotes a slightly faster antigen-positive relapse. This behavior is more intense for SC_2_, even improving CAR-T cell expansion and forming the memory pool for P22. In this scenario, the low cytotoxicity *g*_0_ allows more tumor cells to survive the CAR-T expansion phase, ultimately affecting their depletion. In summary, a smaller value of *d*_*P*_, representing a more homogeneous population of Ag+ cells, accelerates the antigen-negative relapse and delays the antigen-positive relapse.

### 3.4 CAR-T cell cytotoxicity and tumor evolution in relapsed patients

In this section, we investigate the impact of CAR-T cell cytotoxicity on tumor evolution in relapsed patients. Keeping in mind that scenarios SC_1_ and SC_2_ assume different values for *θ, g*_0_ and *T*_*N*0_, we selected patients who share the same parameter values between scenarios, except for *γ* and *a*, which are parameters related to CAR-T cell killing mechanisms. As a result, we conducted a comparative analysis of the cytotoxicity function for patients P12 and L3 from the *Relapse*+ cohort and patients M44 and P22 from the *Relapse*− cohort. Despite the fact that Ag+ tumor cells of patients with positive relapse lose less antigen than in those with negative relapse, Ag+ tumor cells avoid CAR-T antitumor activity due to inhibitory signaling. Cancer cells in patients from the *Relapse*− cohort, on the other hand, bypass CAR-T activity by evading antigen binding.

The evolution of the total tumor cells over time is shown in the upper panel of Figure 5. Until CAR-T cells reach their peak time (*C*_*peak*_), mainly Ag+ tumor cells are depleted and antigen density is downregulated, leading to resistance. When Ag+ tumor cells are exhibiting a resistant phenotype, CAR-T cell cytotoxicity is impaired, creating an opportunity for tumor re-growth and possible escape. However, as CAR-T cell numbers decrease and time approaches *T*_*min*_ (time tumor has reached its minimum population) the therapeutic pressure decreases, leading tumor cells to restore antigen presentation, which enables even further depletion until *T*_*min*_, as can be observed in the heatmaps of Figure 5. Since we set the minimum fraction of CAR-T cell cytotoxicity, *g*_0_, at very low levels for all patients in SC_2_, this phenomenon is well marked with a less pronounced tumor cell decay than in the SC_1_.

**Figure 5:**
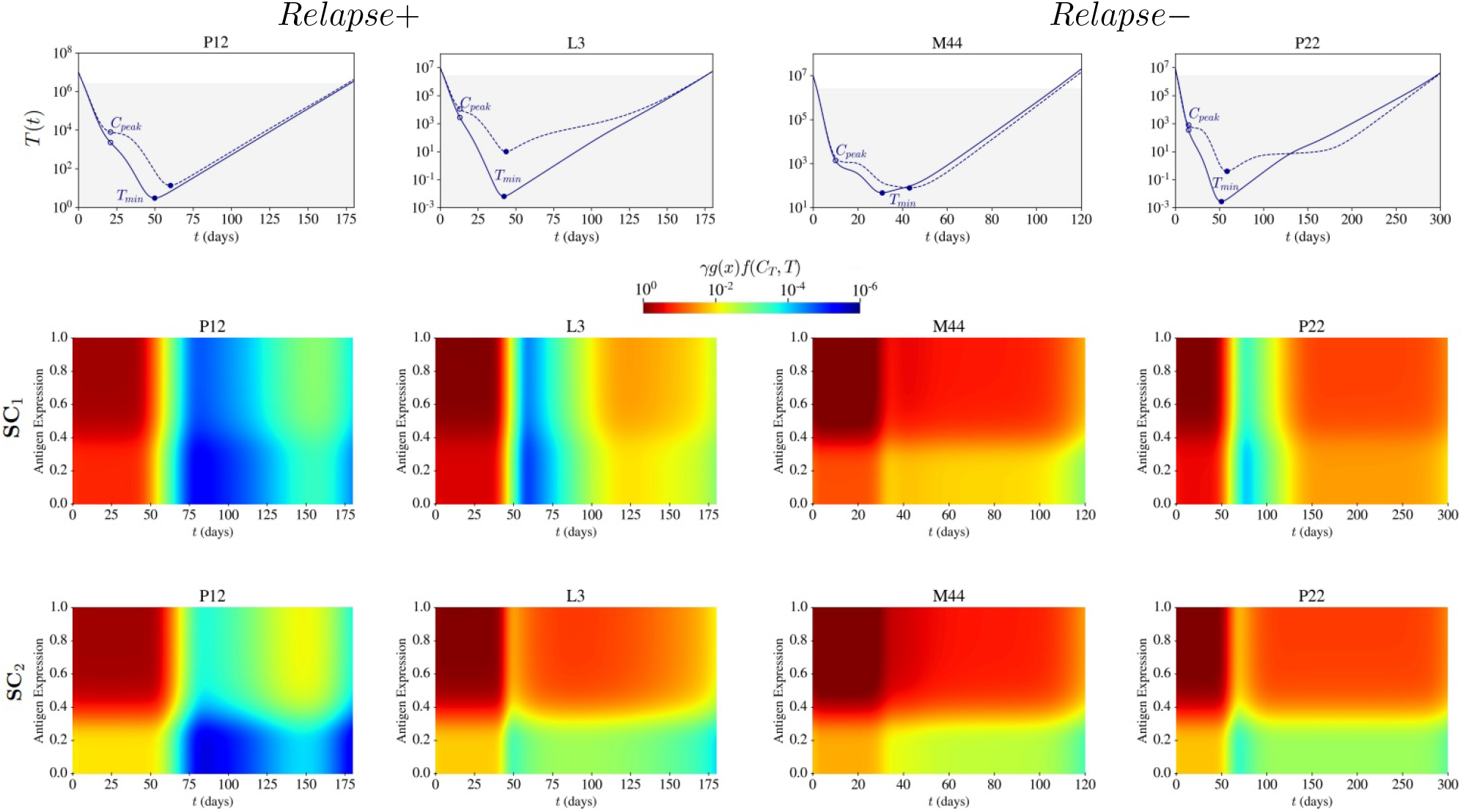
Different therapy scenarios assume different CAR-T cell cytotoxicity, resulting in different tumor dynamics. The upper panel shows the time evolution of the total tumor cells *T* (*t*) for scenario 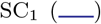 and scenario SC_2_ (). The rapid depletion of tumor cells that occurs at the beginning of CAR-T cell therapy (until *C*_*peak*_) is hindered due to antigen loss. As the CAR-T cell population contracts and the therapy pressure decreases, antigen expression is restored, and the remaining CAR-T cells further reduce the tumor population (until *T*_*min*_. However, if cytotoxicity is insufficient (*Relapse*+) antigen-positive tumor cells escape; and if there is permanent antigen loss (*Relapse*−), the CAR-T cells cannot recognize its target, allowing the antigen-negative tumor cells to escape, although there is a high cytotoxic activity against antigen-positive cells. The middle and bottom panels display the cytotoxicity heatmaps, *γg*(*x*)*f* (*C*_*T*_, *T*), for SC_1_ and SC_2_, respectively. For patients *Relapse*+, CAR-T cytotoxicity is severely impaired due to the minimal number of effector cells in the system and their weak antitumor activity, which cannot avoid antigen-positive cells escape. In patients, *Relapse*−, the mechanism of escape is related to the loss of the target-antigen and not due to CAR-T cytotoxicity. The points *C*_*peak*_ and *T*_*min*_ represent the time when the CAR-T cells reach its peak and when the tumor has reached its minimum population. All parameters used in this simulation are in Table SM-4.

During tumor escape, the heatmaps in Figure 5 show that patients with negative relapse experience a decrease in cytotoxicity, which is partially restored due to the presence of some effector CAR-T cells in the system, which is clearly visible in patient P22. On the other hand, for *Relapse*+ patients, there is a permanent reduction in CAR-T cell cytotoxicity.

### 3.5 Mutation and selection of resistant cells

Now we investigate the fundamental differences between the proposed scenarios. On one hand, we hypothesized that either antigen-negative cells may arise due to mutations (SC_2_) or they were already present before treatment (SC_1_). On the other hand, in our model, these cells can be eliminated by antigen-independent mechanisms, which is defined by the rate *γg*_0_.

We associated the mutation rate *θ* with the progression-free survival (PFS) day for relapsed patients and with the last follow-up day for CR patients. Figure 6a shows that the mutation rate *θ* is negatively correlated with time in both groups. This means that the lower the mutation rate, the longer is the response time. Overall, patients with Ag− relapse showed much greater mutation rates than CR patients. Among the patients in the *Relapse*− cohort, M19, and P22 showed a lineage switch (LS), while B10 and B18 presented early relapse, indicating a higher mutation rate compared to other patients in the *Relapse*+ cohort.

**Figure 6:**
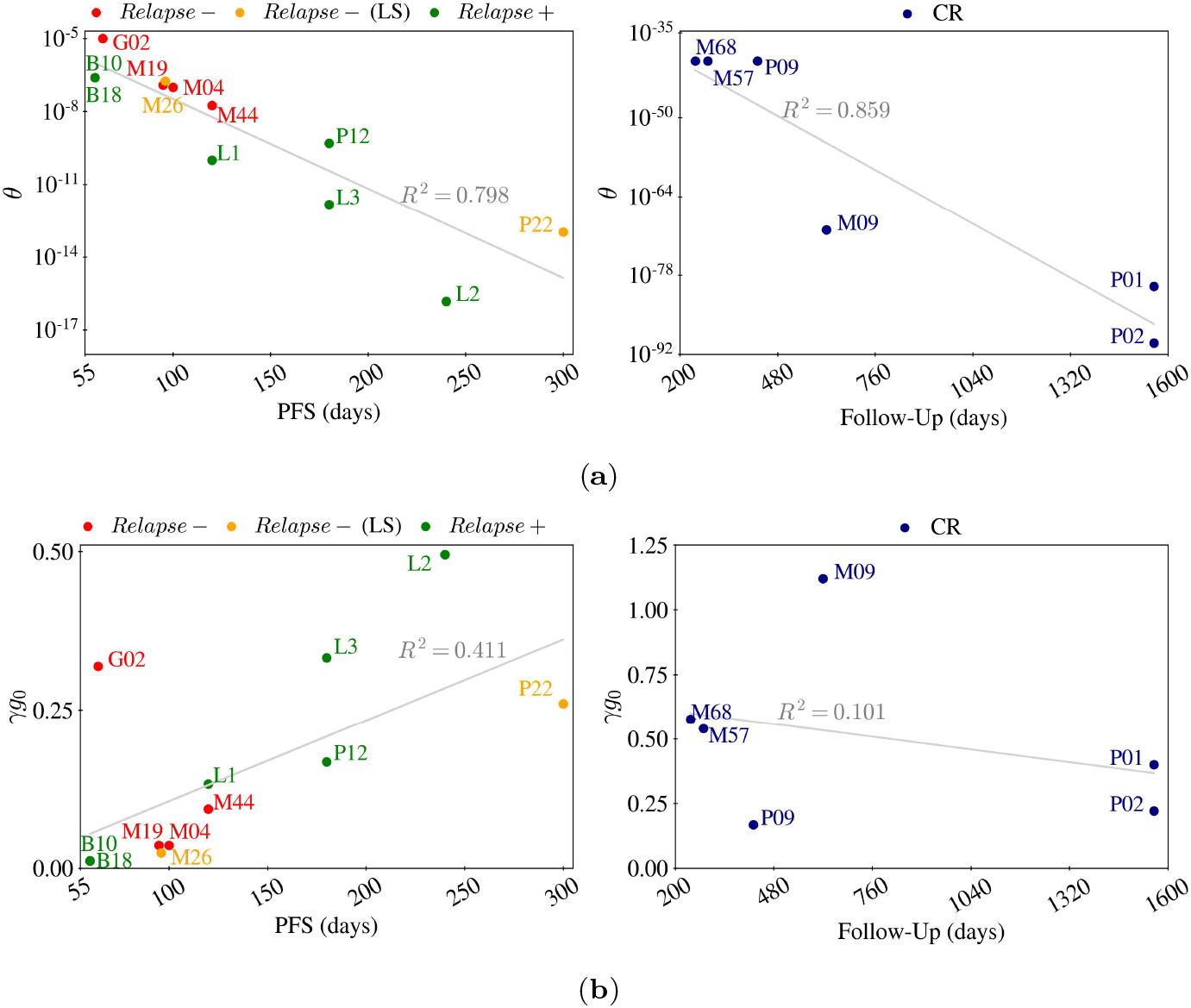
Scatter plots of the fundamental parameters underlying the scenarios proposed: SC_1_ (no mutation, pre-existing Ag− cells) and SC_2_ (onset of Ag− from mutation after therapy start). LS correspond to *Relapse*− that undergo lineage switch. (a) Mutation rates (*θ*) and PFS or last follow-up day for each patient in SC_2_. There is a negative correlation between mutation rate *θ* and response duration, with *Relapse*− patients exhibiting greater mutation rates, and CR patients displaying the lowest rates. Patient G02 was excluded from this analysis because the initial antigen distribution data was already available so we did not consider the scenario where there would be mutations in addition to the initial resistant cells. (b) Antigen-independent killing mechanisms (*γg*_0_) and PFS or last follow-up day considering SC_1_. There is a positive correlation between cytotoxicity and response duration for relapsed patients while CR patients display a weak correlation. PFS, progression-free survival.

The CAR-T cytotoxicity upon Ag− cells (*γg*_0_) was assessed only for SC_1_, as *g*_0_ was the same for all patients in SC_2_. Figure 6b shows that cytotoxicity has a positive correlation with the PFS day in relapsed patients. The highest rates of cytotoxicity for the *Relapse*− group were seen in patients with either a significant initial burden of resistant cells (G02) or a late relapse with lineage switch (P22).

Patients with late relapse in the *Relapse*+ and CR groups also showed significant antigen-independent cytotoxicity, with a weak correlation between *γg*_0_ and response time. This implies that the hypothesis in SC_2_, which assumes the absence of Ag− cancer cells, is more plausible than the one in SC_1_, which would require high antigen-independent CAR-T cell cytotoxicity against these Ag− cells.

## 4 Discussion

In this study, we developed a mathematical model to investigate tumor response to CAR-T cell immunotherapy. Specifically, we focused on understanding the resistance mechanisms which can lead to tumor relapse. To achieve this, we considered two subpopulations of tumor cells characterized by their antigen expression, namely antigen-positive and antigen-negative cells, as well as two phenotypically different populations of CAR-T cells, the memory and effector cells. Our model accurately reproduced the dynamics of tumor and CAR-T cells across various diseases and therapeutic responses. Although the simulations were performed for a specific target antigen (CD19), there are no current limitations to the model’s application to other antigens.

In a more general context of resistance to anti-cancer drugs, Greene et al. [40] proposed a mathematical model that takes into account a continuous variable to describe the cellular “level of resistance” for patients undergoing chemotherapy. Building on this concept, Álvarez-Arenas et al. [41] analyzed a heterogeneous tumor population, including both sensible and resistant cells, with phenotypic plasticity to evaluate the effects of various resistance mechanisms, including Darwinian selection, Lamarckian induction, and nonlocal transfer of extracellular microvesicles on tumor evolution. However, regarding CAR-T cell immunotherapy, so far we are only aware of the work of Liu et al. [35], which proposed a model capable of predicting antigen-negative relapses. Unlike our model, it does not consider the level of antigen expression of tumor cells during the dynamics. Through the incorporation of antigen density as an independent variable, we were able to describe different biological mechanisms of resistance, such as trogocytosis, post-transcriptional antigen downregulation and mutations. By doing so, we provided insights into the role of each resistance mechanism along the dynamics.

Most of the available therapy data indicate the total population of CAR-T cells, while the number of tumor blasts is often expressed as a percentage of nucleated marrow cells. However, there is a particular interest in describing antigen expression profiles, since they are associated with escape from immunotherapy. For example, Nerreter et al. [39] showed that ultra-low expression levels of CD19 are sufficient for inducing cytolytic activity of CD19 CAR-T, but insufficient for inducing cytokine secretion and proliferation. These features motivated us to establish a threshold for the antigen density to regulate antigen-driven mechanisms. Overall, our simulations showed that patients with positive relapse lose less antigen when compared to patients with negative relapse. These findings reveal that antigen-dependent mechanisms play different roles with respect to therapy outcomes.

Following Hamieh et al. [12] findings, our results show that sensible tumor cells that are not killed during CAR-T expansion are susceptible to transient antigen loss, through trogocytosis and therapy pressure. Additionally, the simulations show that in patients with negative relapse, a slow antigen loss followed by a fast tumor depletion does not necessarily result in durable responses. In this cohort, a larger number of effector CAR-T cells are re-stimulated, leading to continuous tumor killing, which ultimately helps in tumor control. On the other hand, in patients with positive relapse, the same mechanism opens up an escape window, likely due to the different CAR-T kinetics observed in this group. It is well-known that in this type of relapse, CAR-T cells exhibit poor cytotoxicity and persistence, which means that even if tumor cells are sensible to treatment, a low level of CAR-T cells is insufficient to control tumor growth. A better understanding of the different mechanisms involved in the dynamics can contribute to the development of combined therapies, new CAR designs and infusion strategies.

Antigen binding plays a pivotal role in tumor clearance. Li et al. [50] showed that upon CAR-T cell therapy the elimination of glioma cells was not immediate and multiple CAR T-cells can interact with these tumor cells. They also showed that the number of CAR T-cells binding to glioma cells depends on the antigen receptor density levels. Indeed, our model shows that, under antigen down-regulation, tumor cells bypass CAR-T cell binding, and tumor depletion is impaired. Our results showed that antigen-mediated CAR-T cell cytotoxicity was effective in preventing the escape of sen-sible cells in *Relapse*− patients, but could not control the emergence of resistant clones. Targeting multiple antigens and increasing antigen presentation and recognition are strategies to overcome this mechanism. In *Relapse*+ patients, cell-to-cell interactions and other antigen-independent mechanisms slowed the growth of antigen-negative cells, although they were not fully eliminated. Nevertheless, in these patients, after CAR-T cell contraction, the low levels of effector cells in the system were insufficient to re-ignite an antitumor response. Enhancing CAR-T cell persistence with a different CAR structure or re-infusing CAR-T cells may be alternatives to surpass this type of escape.

In their review, Lemoine et al. [51] reported that the percentage of relapses due to CD19-loss varies between 7-25% in B-ALL and approximately one-third in DLBCL. In MCL, 23% of relapses were reported in the ZUMA-2 trial, with only one case having undetectable CD19. Moreover, the authors indicate that relapses with a target-Ag−negative clone may occur through immune editing, resulting in the selection of a pre-existing Ag−negative subclone, or possibly through acquired loss of the target-Ag that was initially expressed by the tumor cells. To further explore these hypotheses, we defined two different scenarios: in scenario 1 (SC_1_), we assume that the initial tumor population would have at least 1 antigen-negative cell and that no additional mutations occur; in scenario 2 (SC_2_), resistant cells arise due to mutations and CAR-T cell cytotoxicity against this population are minimum. We considered both scenarios for all relapsed patients to evaluate their feasibility across cohorts. Our findings show that if patients who later presented a positive relapse had one resistant cell at the beginning of the treatment (SC_1_), higher rates of cytotoxicity against antigen-low/-negative tumor cells would be required. Therefore, it seems more likely that these patients’ tumors undergo mutations. For patients with negative relapse, this question is still open. Since detecting such a small number of antigen-negative cells is challenging, strategies like targeting a different antigen or using CAR-T cells as a bridging therapy should be considered.

The development of mathematical models for CAR-T cell immunotherapy remains challenging since many mechanisms are still not fully understood. In particular, some mechanisms such as trogocytosis have only been observed in mice, while data on antigen-density profiles and tumor cell counts are often unavailable. One limitation of our model is that it does not account for CAR-T cell exhaustion, which can affect the number of effector cells in the system and modify antigen modulation, especially in the context of therapy-induced antigen downregulation. As demonstrated by Paixão et al. [25], calculating the fraction of non-exhausted CAR-T cells has the potential to serve as a predictive marker for different responses, but its applicability in the case of relapse requires further investigation. Furthermore, incorporating the interplay between CAR-T cells and other endogenous immune cells may help explain how immune mechanisms in the tumor microenvironment influence treatment outcomes [22, 52]. Despite these limitations, our proposed model successfully recapitulated several resistance mechanisms, described antigen modulation dynamics, and generated new insights into CAR-T cell immunotherapy.

## 5 Conclusions

The proposed model accurately reproduced the dynamics of tumors and CAR-T cells in response to immunotherapy, focusing on the resistance mechanisms that lead to relapse. By incorporating the level of antigen expression as a variable, the model provided insights into the role of each resistance mechanism, shedding light on the antigen-modulation dynamics. Understanding the dynamic features of CAR-T cell therapy, such as effector cell persistence, antigen recognition, and cell-to-cell interactions, may provide new information for developing combinatorial targeting strategies, new CAR designs or CAR-T cell dosing. In future work, we aim to explore these resistance mechanisms for other target antigens and to establish a larger virtual cohort of patients that may help to improve our knowledge of heterogeneous treatment responses.

## Data Accessibility

All data used in this study can be downloaded from the cited sources. All information, such as parameter values and code descriptions used to replicate simulations and analysis are available in this work.

## Authors’ Contributions

Conceptualization, D.S.S., E.A.P., L.R.C.B., R.C.A. and A.C.F.; methodology, D.S.S., R.C.A. and A.C.F.; software, D.S.S.; writing—original draft preparation, D.S.S.; writing—review and editing, D.S.S., E.A.P., L.R.C.B., R.C.A. and A.C.F.. All authors have read and agreed to the published version of the manuscript.

