## SUPPLEMENTARY MATERIAL for "Mechanisms of Resistance to CAR-T cell Immunotherapy: Insights from a Mathematical Model"

Daniela S. Santurio[1,*], Emanuelle A. Paixão[2], Luciana R.C. Barros[3], Regina C. Almeida[4], Artur C. Fassoni[5,†]

[1] Institutional Training Program,[2] Graduate Program, and [4] Computational Modeling Department, Laboratório Nacional de Computação Científica, Petrópolis 25651-075, Brazil
[3] Center for Translational Research in Oncology, Instituto do Câncer do Estado de São Paulo, Hospital das Clínicas da Faculdade de Medicina da Universidade de São Paulo, São Paulo 01246-000, Brazil
[5] Institute for Mathematics and Computer Science, Universidade Federal de Itajubá, Itajubá 37500-903, Brazil

### SM-1 Model Assumptions

**A) Infused CAR-T cells ($CD8^+$:$CD4^+$) have an effector phenotype with cytotoxic activity against tumor cells**

During CAR-T cell production, T cells are typically stimulated with cytokines and other growth factors to promote expansion and activation [1]. This process can cause some of the T cells to differentiate into memory cells; however, the amount of memory cells produced varies depending on several factors such as the manufacturing process or the source of T cells used. Furthermore, most clinical trials do not display this data so, to simplify, we assumed that the infused cells have an effector phenotype, $C_T(t)$, enabling them to recognize and destroy antigen-positive tumor cells through cytotoxicity.

**B) Effector CAR-T cells expand in a patient-specific manner after binding on sensible tumor cells**

The expansion of CAR T cells is a complex mechanism that involves various factors, including intrinsic product attributes, such as the types of co-stimulatory domains (such as CD28 or 4-1BB), the origin of the T cells (autologous or allogeneic), as well as recipient-specific characteristics, such as the type of disease and the level of antigen expression of tumor cells. To approach this heterogeneous scenario, we propose that the CAR-T cell expansion is modeled by the $\kappa(t)$ function and the Holling function $T_S(t)/(A + T_S(t))$. Antigen-binding triggers a series of intracellular signaling pathways, leading to the proliferation of the CAR-T cells, as proposed in our previous model [2]. The function $\kappa(t)$ encompasses patient-specific factors that impact the CAR-T cell initial strength ($p_1$), duration ($p_2$), decay ($p_3$), and a baseline ($r_{min}$) level of proliferation. The function $T_S(t)/(A + T_S(t))$ incorporates the fact that CAR-T cell expansion is not unlimited and may be regulated by other biological mechanisms, such as cell division, cellular senescence, and inhibitory signals from the host immune system [3].

**C) Tumor cells, through an antigen-independent mechanism, inhibit effector CAR-T cells**

It is widely accepted that tumor cells present an immunosuppressive environment that limits the effectiveness of immune attack [4], incorporated in our model through the parameter $\alpha$. Membrane-bound ligands, checkpoint receptors, soluble factors, and suppressive immune cell populations (myeloid-derived suppressor cells (MDSCs), tumor-associated macrophages (TAMs), regulatory T cells (Tregs), and cancer-associated fibroblasts (CAFs)) all contribute to the tumor's complex immunosuppressive network [5]. Although these new cells cannot bind to CAR-T cells, they can induce immune escape by avoiding contact or upregulating coinhibitory ligands such as the Programmed Cell Death Ligand 1 (PD-L1) [6].

**D) Effector and memory CAR-T cells present different death rates and cytolytic activity**
For simplicity, we assume that CAR-T cells have only two phenotypes with different characteristics. Effector CAR-T cells, $C_T(t)$, are short-lived and cytotoxic, promoting cytokine release and inflammation during tumor cell attack. To avoid persistent immune response these cells must die or eventually become exhausted. Memory CAR-T cells, $C_M(t)$, have a low apoptosis rate and provide long-term protection. To maintain long-term antigen-specific antitumor activity, a memory pool is formed through a constant differentiation of effector CAR-T cells at a rate $\epsilon$. However, memory cells must be re-exposed to the target antigen to differentiate back into the effector phenotype and activate a secondary response. Therefore, we assume that memory cells have a smaller mortality rate than effector cells ($\mu_M < \mu$) and return to the effector CAR-T cell phenotype at a rate $\theta_M$.

**E) The tumor population constitution is based on their constitutive/genetic level of antigen expression**
In the context of CAR-T cell immunotherapy, resistance is directly correlated to antigen expression. Relapses with a target-antigen (Ag) negative clone may occur either by the selection of a pre-existing Ag-negative subclone or by acquired loss of the target-Ag that was initially expressed by the tumor cells [7]. The eventual presence of antigen-negative tumor cells suggests that these cells can be inherently resistant even in the absence of treatment. Therefore, we incorporated these concepts in our model by defining two genetically distinct tumor populations: one a population of tumor cells that express the target antigen ($T_P$) and another that does not express the target antigen ($T_N$).

**F) Each tumor population ($T_P$ and $T_N$) is represented by a density function which, at time $t$, has an antigen expression $x$, varying continuously on the interval $[0, 1]$**
Relevant features of CAR-T cell biology such as expansion, cytotoxicity and memory formation, are regulated, directly or indirectly, by antigen receptor signaling. We assume that tumor cell populations, $T_P$ and $T_N$, are heterogeneous with respect to the antigen expression $x \in [0, 1]$. This range was defined by normalizing the values of antigen expression proposed by Nerreter et al. [8], who measured the number of molecules per cell using direct stochastic optical reconstruction microscopy (dSTORM). Since the quantification of antigen expression is dependent on the detection limits of the measuring technique used (e.g., flow cytometry (FC) and quantitative immunofluorescence (MFI)), it is important to emphasize that the antigen density distribution limits should be adjusted accordingly. Detailed information on the normalization and re-scaling are provided in Section SM-4.1.

**G) Antigen-positive tumor cells switch phenotypes during therapy**
Antigen-positive tumor cells may experience permanent and temporary antigen loss. Driven by different mechanisms, these two types of antigen loss occur at different time scales. In our model, permanent antigen loss is due to mutations ($\theta$) and occurs at a much slower rate than transitory antigen loss, which is induced by therapy pressure ($k_I$) [9]. Mutations change the constitutive (or genetic) antigen level expressed by tumor cells, while therapy-induced pressure requires the presence of the CAR-T cells and induces changes only in the phenotype of the tumor cells. Therefore, when there are many CAR-T cells in the system, there can be a significant reduction in antigen expression of antigen-positive cells, which can make them temporarily resistant to therapy. However, when CAR-T cells decay, antigen-positive tumor cells restore their antigen expression and return back to their original heterogeneous distribution of sensible phenotype.

**H) Antigen-negative tumor cells are resistant and do not change their antigen expression**
During CAR-T cell immunotherapy, tumor cells downregulate their antigen expressions to avoid recognition and evade the immune response [10]. However, research has shown that tumor cells can eventually restore their basal levels of antigen expression without treatment [11]. We hypothesized that changes in antigen expression are meaningful only for the antigen-positive tumor population, and that the intrinsic antigen downregulation of the antigen-negative tumor cell population promotes resistance to treatment. The survival advantage of negative-clones will already impair treatment efficacy, therefore considering any additional antigen loss would have little effect on the overall dynamics.

**I) Mutation is a one-way mechanism**
We assume that when antigen-positive tumor cells undergo division, they may mutate to a resistant genotype at a rate $\theta$. The decrease in the antigen expression is permanent and is passed down from mother to daughter cells and does not revert back to the original level of expression [12].

**J) Random epigenetic factors yield infinitesimally small phenotypic modifications in antigen-positive tumor cells**
Cancer cells are known to exhibit epigenetic heterogeneity, which is a type of phenotype heterogeneity without genetic variation. The switching between phenotypic states can occur in a spontaneous and stochastic fashion due to gene expression noise [13]. As proposed in [14], this random instability yields infinitesimally small phenotypic changes in antigen expression, modulated by the diffusion coefficient $\sigma$. Detailed information on this parameter estimation can be found in Section SM-4.2.

**K) Antigen-positive tumor cells restore their antigen expression**
Through a process called trogocytosis, Hamieh et al. showed that for mouse models, CAR-T cells remove target antigens from tumor cells and internalize them, leading to decreased antigen density in cancer cells [11]. However, tumor cells are able to restore antigen expression after a short-term culture without CAR-T cells. Assuming that these mechanisms also lead to variation in antigen expression in human tumor cells, we incorporated the so-called phenotypic drift velocity function, $v(x,t)$, into our model. This function represents a tumor cell's ability to regain antigen and maintain homeostasis in the absence of any external stimulus, such as those due to therapy. Detailed information on the estimation of the velocity rate that modulates this phenotypic drift can be found in Section SM-4.3.

**L) Non-Mutational mechanisms may drive reversible antigen loss**
Experimental evidence shows that antigen negativity followed by CAR-T cell therapy cannot be reduced to a mutational cause. On the opposite, cancer cells may use a broad array of mechanisms to achieve an antigen-negative phenotype [15]. According to Fry et al. [16], 8 of the 12 patients experienced a relapse within a year post CD22-CAR infusion. Interestingly, among these 8 patients, 7 did not experience a complete loss of antigen but instead showed a decrease in CD22 surface expression. This decrease was not due to any mutations in CD22 or changes in mRNA levels. Sotillo et al. [17], found a new mechanism that removes the target epitope from the cell surface without discarding the entire target protein, which does not trigger killing by CD19 CAR-T cells at physiologic levels. These data suggest that antigen expression on the surfaces of cells may be regulated post-transcriptionally, post-translationally, or through the creation of alternative isoforms. Thus, to incorporate such a mechanism into the model, we assembled the function $\bar{x}(t)$ that uses the half-saturation constant $k_I$ to regulate the phenotypic transition due to treatment pressure. Detailed information on $k_I$ estimation can be found in Section SM-4.4.

**M) CAR-T killing incorporates several biological factors.**
CAR-T cell killing is a complex and multi-modal mechanism. Upon antigen-binding, CAR-T cells form a non-classical immune synapse and mediate their anti-tumoral effects through the release of perforins and granzymes to lyse tumor cells, as well as cytokines to sensitize the tumor stroma [18]. Tumor clearance is highly dependent by both patient-specific characteristiscs and CAR design. Majzner et al. [19] demonstrated that the efficacy of CAR-T cells targeting CD19 or HER2 is proportional to target antigen density, but that CD28 endodomain-containing CARs outperform 4–1BB endodomain-containing CARs in response to targets with low antigen density. Pillis et al. [20] used both *in vitro* and *in vivo* experiments to indicate that cytotoxic T cells follow a fractional-kill law with a saturation level, implying that there exists an upper limit at which T cells can interact and kill tumor cells. Li et al. [21] showed two different aspects of glioma cell killing by CAR-T cells. First, the elimination process of glioma by CAR-T cells is not immediate, and second, multiple CAR-T cells can interact with glioma cells depending on the antigen receptor density levels. Altogether, we assumed that CAR-T cells have a maximum cytotoxic rate of $\gamma$, that is antigen-regulated by $g(x)$ and saturated by $f(C_T, T)$.

### SM-2 Clinical Data

To qualitatively characterize clinically important dynamics ranging from complete long-term response to transient response followed by CD19+/CD19- relapse, we collected data from patients treated with anti-CD19 CAR-T cell therapy [22, 23, 24, 25, 26]. Although some studies disclose raw data, for others we obtained data using WebPlotDigitizer [27] from the published figures.

The first dataset [22] is a phase I study with pediatric B-ALL patients using a second-generation CAR with a 4-1BB intracellular costimulatory domain. All patients have measurements of CAR-T cell counts (cell/$\mu$l) during the first month and, patients 04 and 44 were further examined, with data until 3 and 4 months, respectively. We selected patients 09 and 57 who had a complete response (lasting more than 8 months), and patients 04,19,26, and 44 who had a CD19-negative relapse. For this final group of patients, we used the progression-free survival (PFS) days to indicate the analysis duration, from which we set the relapse day with the absolute number of tumor cells greater than the qPCR detection limit.

We also used data of patient 2 from [24], who received an infusion of $1.4\times10^6$/kg CAR-T cells, corresponding to a total dose of $1.08\times10^7$ cells given to a 10-year-old girl weighting approximately 40 kg. CD19 expression analysis in bone marrow samples revealed that the pretreatment blast population is made up of approximately 7.7% CD19- cells and 92.3% CD19+ cells. Due to the lack of measurements in the peripheral blood, we assumed the same percentage of antigen-negative and positive cells to set the initial tumor load. The patient had a clinical relapse that was apparent in the peripheral blood two months after infusion, as evidenced by the reappearance of blast cells in the circulation. These cells were CD45+dim, CD34+, and did not express CD19.

In [23] raw data was provided for adult patients with CLL treated with autologous T-cells transduced with a CD19-directed CAR. Patient outcomes at the last recorded follow-up were available and we selected patients 01,02 and 09 who presented long-term complete remissions (CR), CD19+ relapse, and a transformed CD19-dim DLBCL, respectively.

We used data of patients 10 and 18 from [25]. Both patients had a CD19+ relapse, after receiving allogeneic CAR-T cells with a CD28 co-stimulatory domain without lymphodepletion chemotherapy. Individual doses were given per kilogram, therefore to estimate the total number of CAR-T cells infused, we assumed that an adult patient weighed 60 kg.

Lastly, from [26], patients 1,2, and 3 were diagnosed with ALL and were treated with autologous CAR with CD28 intracellular co-stimulatory domain. After 4-8 months, relapsed CD19+ leukemia cells were confirmed by flow cytometry.

An overview of patients' characteristics and the infused dose is displayed in Table SM-1. Different clinical trials may require different measuring techniques (flow cytometry or qPCR) with different detection limits. In order to obtain feasible predictions, before using *in silico* models it is critical to define data units, measurements conversions, and set mathematical limits within clinical definitions. All this information is detailed in the next section (Section SM-3).

Table SM-1: Individual CAR-T cell dosage, the end of analysis, disease and outcome.

| Patient | CAR-T dose ($\times 10^6$) | Weight (kg) | Disease | End of analysis (days) | Outcome |
|---|---|---|---|---|---|
| | | | **Ma et al., [22]** | | |
| M04 | 0.73 (cells/kg) | 26 | B-ALL | 99 $^{\dagger}$ | *Relapse−* |
| M09 | 1.00 (cells/kg) | 20.5 | B-ALL | 620$^{\dagger}$ | *CR* |
| M19 | 0.50 (cells/kg) | 17 | B-ALL | 95$^{\dagger}$ | *Relapse−* |
| M26 | 0.50 (cells/kg) | 43 | B-ALL | 96$^{\dagger}$ | *Relapse−* |
| M44 | 0.50 (cells/kg) | 37 | B-ALL | 120$^{\dagger}$ | *Relapse−* |
| M57 | 0.50 (cells/kg) | 54 | B-ALL | 280$^{\dagger}$ | *CR* |
| M68 | 0.50 (cells/kg) | 21 | B-ALL | 244$^{\dagger}$ | *CR* |
| | | | **Grupp et al., [24]** | | |
| G02 | 1.4 (cells/kg) | 40* | ALL | 64$^{\dagger\dagger}$ | *Relapse−* |
| | | | **Porter et al., [23]** | | |
| P01 | 1130.0 (cells) | - | CLL | 1560 | *CR* |
| P02 | 14.2 (cells) | - | CLL | 1560 | *CR* |
| P09 | 170.0 (cells) | - | CLL | 422 | *CR* |
| P12 | 118.0 (cells) | - | CLL | 180 | *Relapse+* |
| P22 | 86.4 (cells) | - | CLL | 300 | *Relapse−* |
| | | | **Brudno et al., [25]** | | |
| B10 | 7.8 (cells/kg) | 60* | MCL | 60 | *Relapse+* |
| B18 | 3.1 (cells/kg) | 60* | DLBCL | 60 | *Relapse+* |
| | | | **Li et al., [26]** | | |
| L1 | 13.3 (cells) | - | ALL | 120$^{\dagger\dagger}$ | *Relapse+* |
| L2 | 199.6 (cells) | - | ALL | 240$^{\dagger\dagger}$ | *Relapse+* |
| L3 | 44.3 (cells) | - | ALL | 180$^{\dagger\dagger}$ | *Relapse+* |

* Estimated based on the patient's age.
† Estimated based on the progression-free survival day.
†† Estimated using WebPlotDigitizer

#### SM-3 Processing clinical data

##### SM-3.1 Tumor burden

In clinical studies, tumor burden (TB) is frequently expressed as the number of blasts as a percentage of nucleated marrow cells (%) and is usually measured prior to lymphodepletion chemotherapy. However, because the proposed model is cellular, the tumor load must be converted into the number of cells. According to Lee et al.[28], the absolute number of circulating blasts in the peripheral blood of B-ALL patients can range from below $10^4$ to $10^{10}$, displaying great heterogeneity even between responders and nonresponders. Given the lack of information on tumor load before the CAR-T infusion and any reliable relationship between cell count and % of TB, we set an intermediate initial tumor load of $10^7$ cells for all patients.

##### SM-3.2 CAR-T cells

The dynamic monitoring of CAR-T cell count has enabled the assessment of cellular kinetic parameters associated with heterogeneous responses and therapeutic efficacy. Clinically, the amount of

CAR-T cells in peripheral blood is frequently measured using quantitative polymerase chain reaction (qPCR), normally in the unit of copies CAR/$\mu$g DNA. From the detection limits used by Lee et al.[28], we can establish:

$$10 \text{ CAR-T copies/100ng DNA} = 10^4 \text{ CAR-T cells (absolute number)},$$
$$1 \text{ CAR-T copy/}\mu\text{g DNA} = 10^5 \text{ CAR-T cells.}$$

##### SM-3.3 Detection limits

Laboratory tools are necessary for patient monitoring and in most clinical trials the main methods for evaluating CAR-T cell counts are quantitative PCR (qPCR) or its phenotypic expression by flow cytometry (FC). The qPCR method results are well correlated with those of flow cytometry, although, the assay appears to be more sensitive than flow cytometry [29]. Nonetheless, both methods have a detection limit and, unless specified, we use the same detection limit specified in [23] either for CAR-T or tumor cells, defined as 25 copies/$\mu$g DNA or 2.5x$10^6$ cells for qPCR.

##### SM-3.4 Converting data from bone marrow to peripheral blood

Data acquired in peripheral blood is typically measured in terms of cells/$\mu$L. Assuming that the average volume of blood sample is 5 mL and that the human body contains 5L of blood [30], the number of cells in the entire peripheral blood can be calculated (1$\mu$L $= 10^{-6}$ L). Moreover, compared to the bone marrow, about 1% of cells are present in peripheral blood at all times [31] leading to:

$$1 \text{ cell/}\mu\text{L} = 5\text{x}10^8 \text{ cells .}$$

##### SM-3.5 Defining clinical outcomes

Response assessment was taken at different time courses following CAR-T cell infusion. Patients who have a complete response (CR) or a partial response (PR) are considered responders. CR is achieved when the tumor is clinically undetectable, and (PR) is defined by a tumor burden that is greater than the detection threshold but less than 50% of the initial tumor burden. Patients with lack of response or nonresponders have either stable disease (SD) or progressive disease (PD). SD is defined when tumor load is between 0.5 and 1.5x its initial size, and (PD) is defined when the tumor is above the last limit. For relapsed patients without a explicit classification we considered that the clinical outcome should be between PR or PD responses.

#### SM-4 Parameter Estimation

The model was adjusted for each individual patient based on the reported clinical data and therapy outcome. Used as a constant for all patients, we show below how parameters $x_P, x_N, d_P, d_N$, and $k_P$ were determined using data from assays, $k_I$ was estimated based on biological assumptions, and $d$ was taken from the literature [32, 2]. We set $n = 8$ for all patients due to the lack of data. The remaining parameters were determined through extensive tests on each patient, resulting in a full set of parameter values for the two proposed scenarios, as shown in Table SM-4.

Data from the considered experimental assays were extracted from the published figures using the WebPlotDigitizer tool [27]. The average values for the experimentally determined parameters were found using a nonlinear least squares method and optimized in Python with the Scipy library's curve.fit algorithm [33], which minimizes the difference between the model and the input data using the Levenberg-Marquardt method. The only exception is $k_I$, that was calculated as shown in SM-4.4.

##### SM-4.1 Homeostatic mean and intrinsic variability of antigen expression

To estimate the homeostatic means, $x_P, x_N$, and the intrinsic variability of antigen expression $d_P, d_N$ of tumor cells we used data from [8]. In this study, Nerreter et al. used a single molecule-sensitive direct stochastic optical reconstruction microscopy (dSTORM) to generate expression profiles of CD19 on myeloma cells and to assess elimination of CD19-positive cells by anti-CD19 CAR-T cells *in vitro*. Density distributions were divided into a CD19-positive subpopulation (CD19+) and a CD19-negative subpopulation (CD19-). We used data of CD19 expression (without CAR-T cells) from patients M012, M016, M019, and M22 provided as logarithmic numbers (natural logarithm, Ln) of molecules per $\mu$m$^2$. The range [-6,3] Ln$\mu$m$^{-2}$ of antigen expression was normalized using

$$x_{norm} = \frac{x - x_{min}}{x_{max} - x_{min}}$$

and the fitted curves are displayed in Figure SM-1.

The homeostatic means $x_P$ and $x_N$ and the intrinsic variabilities (standard deviations of normal distributions) of antigen expression $d_P$ and $d_N$ for both CD19+ and a CD19- subpopulations are displayed in Table SM-2. Due to the lack of such data for the patients used in the simulations and for simplicity, we used the average value for all patients . Specifically, we set: $x_N = 0.2992$, $x_P = 0.6113$, $d_N = 0.0608$ and $d_P = 0.060$.

Table SM-2: Mean and intrinsic variability values of antigen expression for the selected patients using data from [8].

| Patient | CD19+ | CD19- |
|---|---|---|
| M012 | $x_N = 0.33514 \pm 0.00062$ | $x_P = 0.65961 \pm 0.00032$ |
| | $d_N = 0.05655 \pm 0.00065$ | $d_P = 0.06656 \pm 0.00036$ |
| M016 | $x_N = 0.31598 \pm 0.00033$ | $x_P = 0.61654 \pm 0.00128$ |
| | $d_N = 0.05546 \pm 0.00037$ | $d_P = 0.05609 \pm 0.00129$ |
| M019 | $x_N = 0.25052 \pm 0.00112$ | $x_P = 0.45985 \pm 0.00028$ |
| | $d_N = 0.05394 \pm 0.00124$ | $d_P = 0.06847 \pm 0.00033$ |
| M022 | $x_N = 0.29506 \pm 0.00048$ | $x_P = 0.70915 \pm 0.00582$ |
| | $d_N = 0.07721 \pm 0.00056$ | $d_P = 0.04800 \pm 0.00581$ |
| Average | $x_N = 0.29918 \pm 0.00070$ | $x_P = 0.61129 \pm 0.00299$ |
| | $d_N = 0.06079 \pm 0.00078$ | $d_P = 0.05978 \pm 0.00299$ |

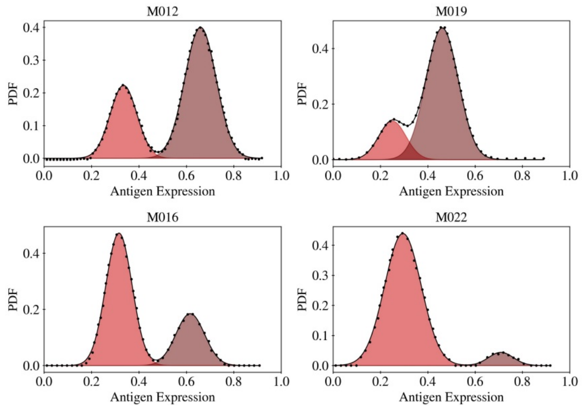


Figure SM-1: Fitted curves for patients M012, M016, M019, and M022. Black dots (•) indicate the extracted data, black line (—) is the fitted curve, the shaded areas, (■) and (■), represent the CD19+ and CD19- density distributions, respectively.

##### SM-4.2 Diffusion coefficient

In our model, the diffusion coefficient is treated as a constant and modulates the epigenetic instabilities in antigen expression that tumor cells exhibit. To estimate its value, we consider that the pre-treatment dynamics is described by the advection-diffusion equation of antigen-positive tumor cells

$$\frac{\partial T_P(x,t)}{\partial t} + \frac{\partial (v(x,t)T_P(x,t))}{\partial x} = \sigma \frac{\partial^2 T_P(x,t)}{\partial x^2} \,, \tag{1}$$

for which $v(x,t) = -k_P(x - x_P)$ in the absence of treatment. Using an unbounded domain for simplicity, with boundarty conditions $T_P(\pm\infty, t) = \dfrac{\partial T_P(\pm\infty, t)}{\partial x} = 0$ and initial condition $T_P(0,t) = u_0(x)$, the steady state solution for (1) is:

$$T_P^{ss}(x) = \frac{A_0}{\sqrt{2\pi}\sqrt{\sigma/k_P}} \exp\left(-\frac{1}{2}\left(\frac{x - x_P}{\sqrt{\sigma/k_P}}\right)^2\right) ,$$

where $A_0 = \int_{\mathbb{R}} u_0(x)$. This shows that, before treatment, the distribution of tumor cells along the antigen expression space $x$ is given by a Gaussian distribution with mean $x_P$ and standard deviation $\sqrt{\sigma/k_P}$, which agree with the data obtained in [8]. Comparing to the data shown above, we conclude that $\sigma$ and $k_P$ should satisfy $\sqrt{\sigma/k_P} = d_P$. We thus set $\sigma = k_P d_p^2$ and estimate $k_P$ as described below in Section SM-4.3)

#### SM-4.3 Antigen density rate of phenotypic transition

To estimate the rate at which tumor cells restore the homeostatic antigen expression, we use data from Hamieh et al. [11]. Using flow cytometry, they measured CD19 expression in NALM6 (ALL) cells retrieved from mice treated with 19-BB$\zeta$ CAR-T cells. We estimated that the CD19 expression at day 0 and day 6 in an *ex vivo* culture was 6.500mol/cell and 11.000mol/cell, respectively. The homeostatic level of CD19 expression was assumed 22.000mol/cell from NALM6 cells measured *in vitro*.

Disregarding tumor proliferation (short-term culture) and any effects caused by CAR-T cells, data shows that from day 0 to day 6 antigen expression was restored from 29.54% to 50% of the homeostatic level. Using our previously calibrated data, we set the homeostatic level as $x_P = 0.6113$ and the levels as $x_P = 0.4307$ and $x_P = 0.3057$ at days 0 and 6, respectively. Figure SM-2 shows a schematic description of the calibratiom process for estimating the homeostatic level $x_P$.

To approximate $k_P$ we solved Equation (1) for $x \in [0, 1]$ and considering zero-flux boundary conditions with the following initial and final conditions:

$$T_P(x, t = 0) = \mathcal{N}(x_P = 0.4307, d_P^2) \qquad \text{and} \qquad T_P(x, t = 6) = \mathcal{N}(x_P = 0.3057, d_P^2)\ , \tag{2}$$

where $d_P = 0.060$, $v(x, t) = -k_P(x - x_P)$ and $\sigma = k_P d_P^2$. Altogether, we found $k_P = 0.095\text{day}^{-1}$.

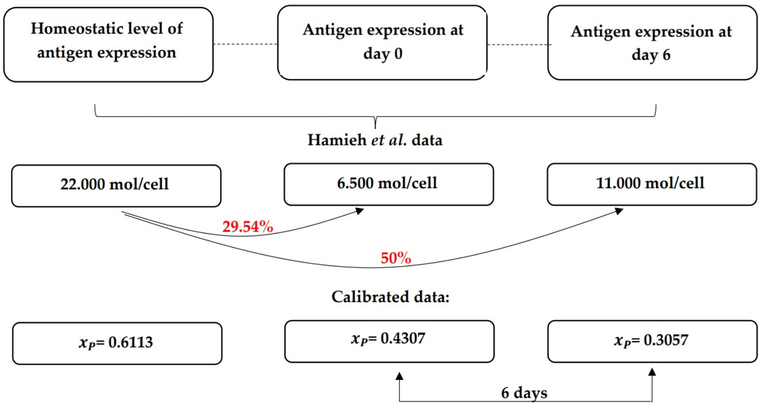


Figure SM-2: Schematic description of the calibration process. From Hamieh et al. [11] data we considered that the homeostatic CD19 expression was 22.000mol/cell. In an *ex vivo* culture, CD19 expression at day 0 and day 6 were 6.500mol/cell and 11.000mol/cell, representing a variation of 29.54% to 50% of the homeostatic level, respectively. Considering the same variation but using our previously calibrated data, we set the antigen expression levels on day 0 and 6 to estimate the drift velocity rate $k_P = 0.095\text{day}^{-1}$.

#### SM-4.4 Antigen modulation by CAR-T cell therapy

Resistant cells may arise during CAR-T cell therapy either through mutations or active, therapeutically induced, non-genetic reprogramming mechanisms. The model incorporates the half-saturation constant $k_I$, that modulates the therapy pressure by regulating the phenotypic transition of the therapy driven level ($\bar{x}(t)$) of antigen expression in $T_P$ (positive) tumor cells. Higher $k_I$ values slow antigen regulation, suggesting tumor cells less susceptible to non-genetic reprogramming events, while lower values lead to faster changes in antigen density. This relationship is illustrated in Figure SM-3.

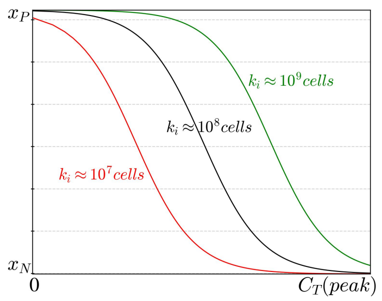


Figure SM-3: Active, therapeutically induced, non-genetic reprogramming mechanisms of antigen regulation. Increasing the value of $k_I$ leads to a slower regulation of antigens, implying that tumor cells become less susceptible to non-genetic reprogramming events.

We assume that before CAR-T cell therapy, tumor cells display their homeostatic level of antigen expression $x_P$. When CAR-T cells reach their peak ($C_T(peak)$), the therapy pressure is at its maximum and tumor cells display their lower level of antigen expression, $x_N$. As a result, we can estimate $k_I$ as follows:

Minimum stress: $\bar{x}(t) = x_P - \delta$ and $C_T(t) = C_T(0)$, therefore

$$x_P - \delta = x_P - (x_P - x_N)\frac{C_T(0)}{k_I + C_T(0)} \Longrightarrow \frac{\delta}{x_P - x_N} = \frac{C_T(0)}{k_I + C_T(0)}$$

Maximum stress: $\bar{x}(t) = x_N + \delta$ and $C_T(t) = C_T(peak)$, therefore

$$x_N + \delta = x_P - (x_P - x_N)\frac{C_T(peak)}{k_I + C_T(peak)} \Longrightarrow \frac{\delta}{x_P - x_N} = 1 - \frac{C_T(peak)}{k_I + C_T(peak)}$$

Combining the above conditions we obtain

$$1 - \frac{C_T(peak)}{k_I + C_T(peak)} = \frac{C_T(0)}{k_I + C_T(0)} \Longrightarrow k_I = \sqrt{C_T(peak)C_T(0)}$$

Table SM-3 displays the individual values of $k_I$ estimated from the extracted data of the selected patients. After the calibration of the constant parameters we were able to perform tests on each patient and establish a full set of parameter values for the two proposed scenarios, as shown in Table SM-4.

Table SM-3: CAR-T cell data and half-saturation constant $k_I$.

| **Patient** | $C_T(0)$ (cells) | $C_T(peak)$ (cells) | $k_I$ (cells) |
|---|---|---|---|
| **Ma et al., [22]** | | | |
| M04 | $1.90 \times 10^{7}$ | $8.35 \times 10^{10}$ | $1.26 \times 10^{9}$ |
| M09 | $2.05 \times 10^{7}$ | $7.17 \times 10^{10}$ | $1.21 \times 10^{9}$ |
| M19 | $8.50 \times 10^{6}$ | $2.95 \times 10^{10}$ | $5.01 \times 10^{8}$ |
| M26 | $2.15 \times 10^{7}$ | $3.60 \times 10^{10}$ | $8.79 \times 10^{8}$ |
| M44 | $1.85 \times 10^{7}$ | $1.38 \times 10^{10}$ | $5.06 \times 10^{8}$ |
| M57 | $2.70 \times 10^{7}$ | $3.33 \times 10^{10}$ | $9.48 \times 10^{8}$ |
| M68 | $1.05 \times 10^{7}$ | $8.14 \times 10^{10}$ | $9.25 \times 10^{8}$ |
| **Grupp et al., [24]** | | | |
| G02 | $5.6 \times 10^{7}$ | $3.89 \times 10^{9}$ | $4.67 \times 10^{8}$ |
| **Porter et al., [23]** | | | |
| P01 | $1.13 \times 10^{9}$ | $4.10 \times 10^{10}$ | $6.81 \times 10^{9}$ |
| P02 | $1.42 \times 10^{7}$ | $2.51 \times 10^{9}$ | $1.89 \times 10^{8}$ |
| P09 | $1.70 \times 10^{8}$ | $8.14 \times 10^{9}$ | $1.18 \times 10^{9}$ |
| P12 | $1.18 \times 10^{8}$ | $6.32 \times 10^{9}$ | $8.63 \times 10^{8}$ |
| P22 | $8.64 \times 10^{7}$ | $1.3 \times 10^{1}0$ | $1.06 \times 10^{9}$ |
| **Brudno et al., [25]** | | | |
| B10 | $4.68 \times 10^{8}$ | $2.00 \times 10^{10}$ | $3.06 \times 10^{9}$ |
| B18 | $1.86 \times 10^{8}$ | $2.00 \times 10^{10}$ | $1.93 \times 10^{9}$ |
| **Li et al., [26]** | | | |
| L1 | $1.33 \times 10^{7}$ | $5.58 \times 10^{9}$ | $2.72 \times 10^{8}$ |
| L2 | $2.00 \times 10^{8}$ | $2.00 \times 10^{10}$ | $2.00 \times 10^{9}$ |
| L3 | $4.43 \times 10^{7}$ | $8.87 \times 10^{8}$ | $1.98 \times 10^{8}$ |

Table SM-4: Individual parameter values, in appropriate units (a.u.), used in the simulations. Patients were divided into three cohorts according to the reported response: Complete Response (CR), Positive Relapse (*Relapse*+), and Negative Relapse (*Relapse*−). The peak-day observed in the experimental data is also informed, which was used to determine the parameter $p_2(p_2^{p_3} = 1/(\text{peak-day})^{p_3})$. Parameters whose values were the same for all patients are: $d = 0.305$, $k_P$=0.095, $x_P$=0.6113, $x_N$=0.2992, $d_P$=0.06, $d_N$=0.608, and $n$=8. Since we proposed two different scenarios, the parameters for Scenario 1 are displayed in black, while any parameter that change its value for Scenario 2 is displayed in red.

| Parameters | *Relapse*− | | | | | | *Relapse*+ | | | | | | CR | | | | | |
|---|---|---|---|---|---|---|---|---|---|---|---|---|---|---|---|---|---|---|
| | **M04** | **M19** | **M26** | **M44** | **G02** | **P22** | **B10** | **B18** | **P12** | **L1** | **L2** | **L3** | **P01** | **P02** | **P09** | **M09** | **M57** | **M68** |
| $r$ | 0.19 | 0.19 | 0.19 | 0.18 | 0.18 | 0.11 | 0.14 | 0.14 | 0.11 | 0.19 | 0.16 | 0.16 | 0.1 | 0.1 | 0.1 | 0.19 | 0.19 | 0.19 |
| $\theta$ | 0 | 0 | 0 | 0 | 0 | 0 | 0 | 0 | 0 | 0 | 0 | 0 | 0 | 0 | 0 | 0 | 0 | 0 |
| $\theta$ | 1×10$^{-7}$ | 1.2×10$^{-7}$ | 1.8×10$^{-7}$ | 1.8×10$^{-8}$ | 1×10$^{-5}$ | 1.05×10$^{-13}$ | 2.5×10$^{-7}$ | 2.5×10$^{-7}$ | 5×10$^{-10}$ | 1×10$^{-10}$ | 1.5×10$^{-16}$ | 1.5×10$^{-12}$ | 1×10$^{-80}$ | 1×10$^{-90}$ | ×10$^{-40}$ | 1×10$^{-70}$ | 1×10$^{-40}$ | 1×10$^{-40}$ |
| $A$ | 7.5×10$^{2}$ | 7.5×10$^{2}$ | 5×10$^{3}$ | 4 | 2.5×10$^{2}$ | 150 | 6.5×10$^{5}$ | 6.5×10$^{5}$ | 5×10$^{3}$ | 2.15×10$^{3}$ | 5 | 1.5×10$^{3}$ | 50 | 2×10$^{3}$ | 35 | 50 | 1.5×10$^{4}$ | 5.5×10$^{3}$ |
| $a$ | 1.80×10$^{6}$ | 1.8×10$^{6}$ | 1.8×10$^{6}$ | 1×10$^{6}$ | 1×10$^{3}$ | 1.8×10$^{6}$ | 6×10$^{6}$ | 1×10$^{7}$ | $5 \times 10^5$ ($1 \times 10^5$) | $1 \times 10^5$ ($2 \times 10^5$) | 40 | 1×10$^{2}$ | 0 | 0 | 0 | 0 | 0 | 0 |
| $k_I$ | 1.26×10$^{9}$ | 5.01×10$^{8}$ | 8.8×10$^{8}$ | 5.3×10$^{8}$ | 4.67×10$^{8}$ | 6.93×10$^{8}$ | 3.06×10$^{9}$ | 2.1×10$^{9}$ | 8.63×10$^{8}$ | 2.72×10$^{8}$ | 2×10$^{9}$ | 1.98×10$^{8}$ | 6.81×10$^{9}$ | 1.89×10$^{9}$ | 1.18×10$^{9}$ | 1.21×10$^{8}$ | 9.479×10$^{8}$ | 9.25×10$^{9}$ |
| $\gamma$ | 1.2 | 1.2 | 1.2 | 1.55 | 1.45 | 1 (1.1) | 0.74 | 0.8 | 0.67 | 0.53 (0.61) | 1.1 | 0.95 | 2 | 0.85 | 1.4 | 3.5 | 1.8 | 1.65 |
| $g_0$ | 0.03 | 0.03 | 0.02 | 0.06 | 0.22 | 0.26 | 0.015 | 0.015 | 0.25 | 0.25 | 0.45 | 0.35 | 0.2 | 0.26 | 0.12 | 0.32 | 0.3 | 0.35 |
| $g_0$ | 0.015 | 0.015 | 0.015 | 0.015 | 0.015 | 0.015 | 0.015 | 0.015 | 0.015 | 0.015 | 0.015 | 0.015 | 0.015 | 0.015 | 0.015 | 0.015 | 0.015 | 0.015 |
| $\alpha$ | 1.15×10$^{-7}$ | 1.15×10$^{-7}$ | 1.3×10$^{-7}$ | 3.1×10$^{-7}$ | 8.7×10$^{-7}$ | 3.2×10$^{-7}$ | 9.2×10$^{-7}$ | 8.1×10$^{-7}$ | 3.34×10$^{-7}$ | 9×10$^{-8}$ | 1.3×10$^{-7}$ | 2.5×10$^{-7}$ | 1.5×10$^{-6}$ | $3.71 \times 10^{-7}$ ($3.64 \times 10^{-7}$) | 7×10$^{-7}$ | $1.563 \times 10^{-6}$ ($1.528 \times 10^{-6}$) | $1.189 \times 10^{-6}$ ($1.17 \times 10^{-6}$) | $1.07 \times 10^{-6}$ ($1.051 \times 10^{-6}$) |
| $\mu$ | 0.8245 | 0.842 | 0.542 | 0.8235 | 0.75 | 0.2985 | 0.848 | 0.7 | 0.25 | 0.5985 | 0.6975 | 0.56 | 0.29995 | 0.095 | 0.1225 | 0.74 | 0.34 | 0.735 |
| $r_{min}$ | 0.001 | 0.001 | 0.001 | 0.4 | 0.001 | 0.7 | 0.001 | 0.001 | 0.001 | 0.001 | 0.001 | 0.001 | 0.001 | 0.35 | 0.001 | 0.001 | 0.001 | 0.001 |
| $p_1$ | 1.95 | 1.95 | 1.95 | 1.475 | 3.3 | 0.45 | 5.76 | 5.78 | 1.64 (1.61) | 1.44 | 1.3 | 1.35 | 4.3 | 1 | 1.51 | 4.71 | 4.765 | 6.16 |
| $p_2$ | 3.16×10$^{-19}$ | 1×10$^{-18}$ | 5.01×10$^{-4}$ | 1×10$^{-30}$ | 5.91×10$^{-4}$ | 0.11458 | 3.24×10$^{-32}$ | 4.44×10$^{-26}$ | 0.1187 | $4.355 \times 10^{-7}$ ($6.957 \times 10^{-6}$) | 1.96×10$^{-5}$ | 3.5×10$^{-5}$ | 0.1462 | 1.623×10$^{-2}$ | 0.1312 | 1.2533×10$^{-17}$ | 2.46×10$^{-3}$ | 1.52×10$^{-23}$ |
| $p_3$ | 18.5 | 18 | 3.3 | 30 | 3.1 | 0.8 | 33 | 30 | 0.7 | 5.55 (4.5) | 4 | 4 | 1.75 | 1.2 | 0.75 | 20 | 2.61 | 27 |
| $\epsilon$ | 0.0255 | 0.008 | 0.083 | 0.0115 | 0.05 | 0.0015 | 0.002 | 0.15 | 0.1 | 0.0015 | 0.0025 | 0.04 | 5×10$^{-5}$ | 5×10$^{-3}$ | 2.5×10$^{-3}$ | 0.01 | 0.01 | 0.015 |
| $\theta_M$ | 10$^{-5}$ | 10$^{-6}$ | 10$^{-6}$ | 5×10$^{-6}$ | 10$^{-6}$ | 10$^{-4}$ | 10$^{-9}$ | 10$^{-9}$ | 10$^{-9}$ | 4×10$^{-9}$ | $2.5 \times 10^{-11}$ ($7.25 \times 10^{-11}$) | $5 \times 10^{-10}$ ($1.1 \times 10^{-9}$) | 10$^{-3}$ | 10$^{-3}$ (×10$^{-5}$) | 10$^{-3}$ | 0.01 | 10$^{-3}$ | 10$^{-4}$ |
| $\mu_M$ | 0.015 | 0.01 | ×10$^{-4}$ | 5.5×10$^{-3}$ | 0.11 | 6×10$^{-3}$ | 0.01 | 0.05 | 0.065 | 5×10$^{-3}$ | 8×10$^{-3}$ | 0.02 | ×10$^{-5}$ | 2×10$^{-3}$ | 0.01 | 1.5×10$^{-3}$ | 5×10$^{-3}$ | 1.5×10$^{-3}$ |
| $T_{P0}$ | 10$^{7}$ | 10$^{7}$ | 10$^{7}$ | 10$^{7}$ | 9.23×10$^{6}$ | 10$^{7}$ | 10$^{7}$ | 10$^{7}$ | 10$^{7}$ | 10$^{7}$ | 10$^{7}$ | 10$^{7}$ | 10$^{7}$ | 10$^{7}$ | 10$^{7}$ | 10$^{7}$ | 10$^{7}$ | 10$^{7}$ |
| $T_{N0}$ | 1 (0) | 1 (0) | 1 (0) | 1 (0) | 0.77×10$^{6}$ | 1 (0) | 1 (0) | 1 (0) | 1 (0) | 1 (0) | 1 (0) | 1 (0) | 1 (0) | 1 (0) | 1 (0) | 1 (0) | 1 (0) | 1 (0) |
| $C_T(0)$ | 1.9×10$^{7}$ | 8.5×10$^{6}$ | 2.15×10$^{7}$ | 1.85×10$^{7}$ | 5.6×10$^{7}$ | 8.64×10$^{6}$ | 4.68×10$^{8}$ | 1.86×10$^{8}$ | 1.18×10$^{8}$ | 1.33×10$^{7}$ | 2×10$^{8}$ | 4.43×10$^{7}$ | 1.13×10$^{9}$ | 1.42×10$^{7}$ | 1.7×10$^{8}$ | 2.05×10$^{7}$ | 2.7×10$^{7}$ | 1.05×10$^{7}$ |
| $C_M(0)$ | 0 | 0 | 0 | 0 | 0 | 0 | 0 | 0 | 0 | 0 | 0 | 0 | 0 | 0 | 0 | 0 | 0 | 0 |

#### SM-5 Additional Results

##### SM-5.1 Model assessment for patients with different therapy outcomes for $SC_2$

The estimated parameter values and the corresponding model simulation for $SC_2$ for each patient are presented in Table SM-4 and Figure SM-4, respectively. Since the initial distribution of antigen expression was available for patient G02, we have not hypothesized a different scenario, therefore we maintained all the parameters in both simulations. Although driven by different conditions and mechanisms, model simulations show good agreement with the available data.

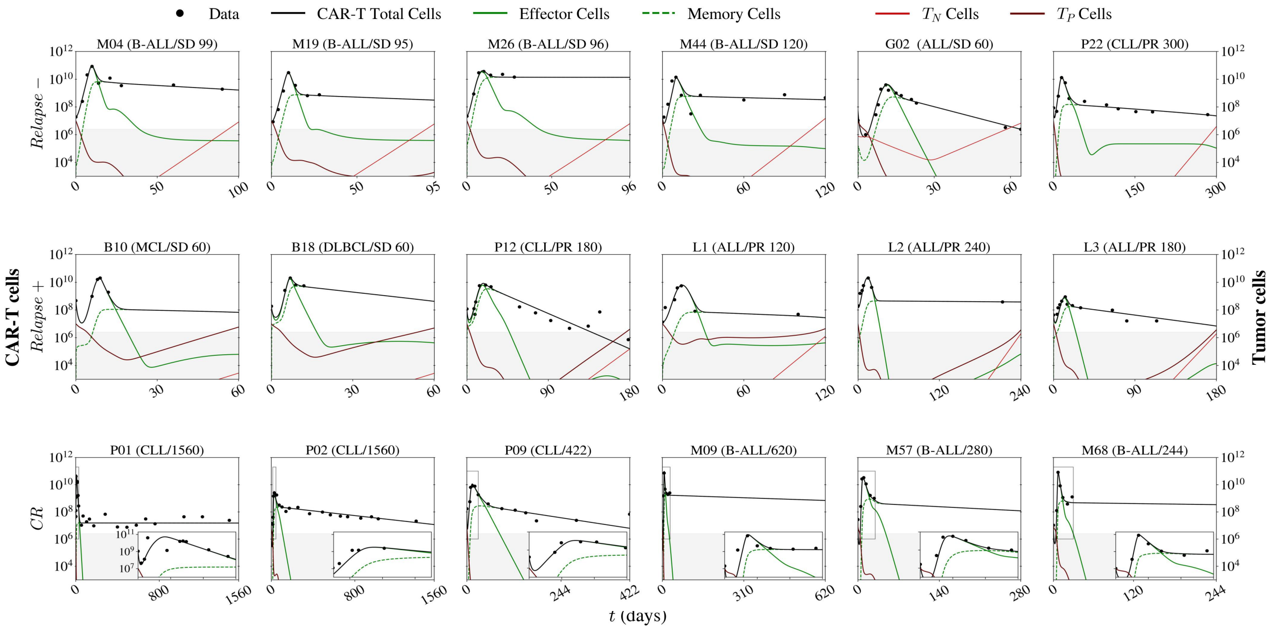


Figure SM-4: Model fits of tumor-CAR-T cell interactions for 18 patients divided into three cohorts: *Relapse*− (top panel), *Relapse*+ (middle panel), and *CR* (bottom panel). The total CAR-T cell population includes effector ($C_T$) and memory ($C_M$) phenotypes, while the total tumor population encompasses antigen-negative ($T_N$) and antigen-positive ($T_P$) cells. The qPCR detection threshold of $2.5 \times 10^6$ cells was represented by the gray area. The zoomed-in analysis of patient P02 was made from day 15 to day 45, while the remaining CR patients were analyzed from day 0 to day 30. B-All, B-cell acute lymphoblastic leukemia; ALL, acute lymphoblastic leukemia; CLL, chronic lymphoblastic leukemia; MCL, mantle cell lymphoma; DLBCL, diffuse large B cell lymphoma. These simulations consider $SC_2$ (Ag- cells arise due to mutations).

##### SM-5.2 Sensitivity Analysis

To perform the sensitivity analysis, virtual patients were created to represent each cohort, including *Relapse*−, *Relapse*+ and CR. Parameter values for each virtual patient were determined using the median (second quartile) values across patients in the same cohort. After ordering the individual values of all six patients presented in Table SM-4 from each cohort, the medians were obtained as the average of the third and fourth values. Figure SM-5 displays the simulations for each virtual patient, and Table SM-5 presents the corresponding parameter values. Notice that the responses of all virtual patient are in accordance with the outcomes of the respective cohort.

Next, we used the Morris method implemented in the SALib library in Python [34] to perform the sensitivity analysis with respect to the state variables. As the quantity of samples necessary to attain convergence for a model with 23 parameters would be unfeasible, we opted to solely analyze the parameters associated with the tumor dynamics and the resistance mechanisms. Perturbations of ± 16.67% were applied to the constant parameters, while we set the range of analysis between the third and fourth values used to generate the virtual patient for the remaining parameters. Figure SM-6 displays the first-order sensitivity index and Table SM-5 presents the range used in the simulations. This global sensitivity index identifies the impact of each analyzed parameter on the quantity of interest. For comparison, the index values were normalized using the sum of all index values in each experiment.

In analyzing the sensitivity index of relapsed (*Relapse*− and *Relapse*+) virtual patients, it is apparent that $\gamma$ and $x_P$ play a significant role in the dynamics of the effector and memory CAR-T cell populations, regardless of the scenario. The former corresponds to the maximum cytotoxic rate of effector CAR-T cells, whereas the latter relates to the homeostatic level of antigen expression in $T_P$ cells. In addition, for the CR patient, the half-saturation constant of the cytotoxic effect on tumor cells, $d$, also appears to affect the CAR-T cell populations. These findings suggest that the cytotoxic strength of effector cells in eliminating tumor cells, as well as the initial level of antigen expression, can modulate CAR-T activity.

Examining the sensitivity index with respect to the antigen-negative tumor cells, it is observed that the parameters $T_{N0}$ and $\theta$ appear as important parameters in $\mathrm{SC}_1$ and $\mathrm{SC}_2$, respectively, mainly in the initial stages of the dynamics. Furthermore, the dynamics of $T_N$ cells are affected by the homeostatic mean levels of antigen expression $x_P$ and $x_N$. While the importance of CAR-T cell cytotoxicity via antigen-independent mechanisms, $g_0$, is evident in the relapsed (*Relapse*− and *Relapse*+) virtual patients, the half-saturation constant of the cytotoxic effect on tumor cells, $d$, and the growth rate of tumor cells, $r$, are important in the CR patient.

When analyzing the antigen-positive tumor cells, it is noteworthy that the CR patient shows a pattern of sensitivity index similar to that of the antigen-negative tumor cells. However, in the *Relapse*− patient, the intrinsic variability of $T_N$ cells, $d_N$, along with the homeostatic mean level of antigen expression $x_P$, strongly impact the behavior of $T_P$ cells, especially during the latter half of the analysis. In contrast, for the *Relapse*+ patient, both $x_P$ and $x_N$ are the parameters that stand out, while the remaining parameters exhibit similar levels of importance.

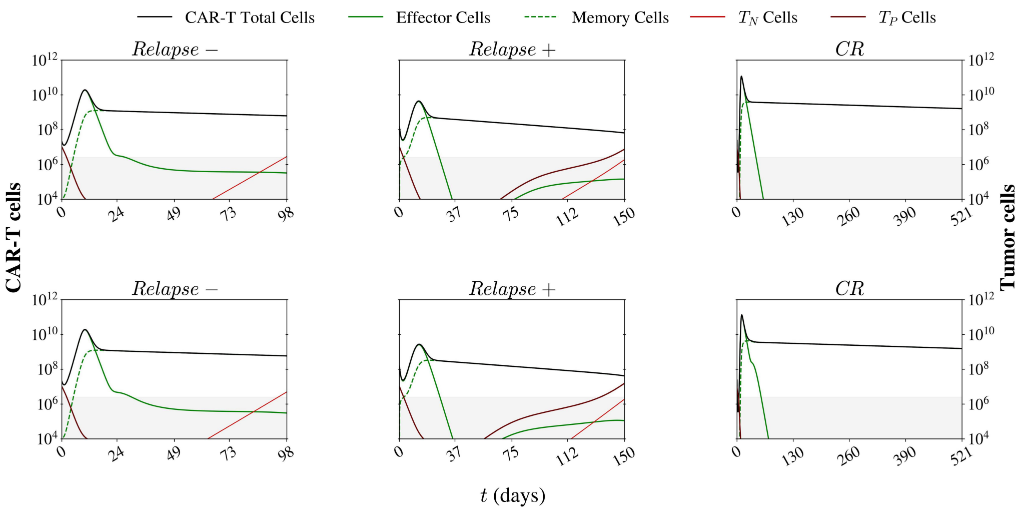


Figure SM-5: *In silico* simulations for Scenario 1 (top panel) and Scenario 2 (bottom panel) of tumor-CAR-T cell interactions for the *Relapse*−, *Relapse*+ and CR virtual patients. The total CAR-T cell population includes effector ($C_T$) and memory ($C_M$) phenotypes, while the total tumor population encompasses antigen-negative ($T_N$) and antigen-positive ($T_P$) cells. The qPCR detection threshold of $2.5 \times 10^6$ cells was represented by the gray area.

Table SM-5: Virtual patients parameter values. Parameters whose values were the same for all patients are $d$ = 0.305, $k_P$=0.095, $x_P$=0.6113, $x_N$=0.2992, $d_P$=0.06, $d_N$=0.608, and $n$=8. Since we proposed two different scenarios, the parameters for Scenario 1 are displayed in black, while any parameter that change its value for Scenario 2 are displayed in red.

| **Parameter** | **Unit** | *Relapse−* | *Relapse+* | *CR* |
|---|---|---|---|---|
| $t$ | day | 98 | 150 | 521 |
| $r$ | day$^{-1}$ | 0.185 | 0.15 | 0.145 |
| $\theta$ | – | 0 | 0 | 0 |
| $\theta$ | – | 1.0 $\times 10^{-7}$ | 3.0 $\times 10^{-10}$ | 5.0 $\times 10^{-41}$ |
| $A$ | cell | $5\times10^2$ | $3.58\times10^3$ | $1.03\times10^3$ |
| $a$ | cell | $1.8\times10^6$ | $3 \times 10^5$ ($1.5 \times 10^5$) | 0 |
| $k_I$ | cell | $6.12\times10^8$ | $1.43\times10^9$ | $1.2\times10^9$ |
| $\gamma$ | day$^{-1}$ | 1.2 | 0.77 | 1.725 |
| $g_0$ | – | 0.045 | 0.25 | 0.25 |
| $g_0$ | – | 0.015 | 0.015 | 0.015 |
| $\alpha$ | (cell.day)$^{-1}$ | $2.2\times10^{-7}$ | $2.92\times10^{-7}$ | $1.13 \times 10^{-6}$ ($1.11 \times 10^{-6}$) |
| $\mu$ | day$^{-1}$ | 0.799 | 0.629 | 0.318 |
| $r_{min}$ | day$^{-1}$ | 0.001 | 0.001 | 0.001 |
| $p_1$ | day$^{-1}$ | 1.95 | 1.54 (1.525) | 4.505 |
| $p_2$ | day$^{-1}$ | $5.32\times10^{-11}$ | $6.82 \times 10^{-10}$ ($3.29 \times 10^{-9}$) | $1.33\times10^{-3}$ |
| $p_3$ | - | 10.65 | 4.775 (4.25) | 2.18 |
| $\epsilon$ | day$^{-1}$ | 0.0185 | 0.02125 | 0.0075 |
| $\theta_M$ | (cell.day)$^{-1}$ | $3 \times 10^{-6}$ ($1 \times 10^{-6}$) | $1\times10^{-9}$ | $1\times10^{-3}$ |
| $\mu_M$ | day$^{-1}$ | 0.008 (0.0085) | 0.015 | 0.00175 |

Table SM-6: Ranges of variation of parameter values used in the sensitivity analysis.

| **Parameter** | *Relapse−* | *Relapse+* | *CR* |
|---|---|---|---|
| $t$ | 98 | 150 | 521 |
| $r$ | [0.18-0.19] | [0.14-0.16] | [0.1-0.19] |
| $\theta$ | 0 | 0 | 0 |
| $\theta$ | [1.0-1.2 $\times10^{-7}$] | [1.0 $\times10^{-10}$-5.0 $\times10^{-10}$] | [1.0 $\times10^{-70}$-1.0 $\times10^{-40}$] |
| $a$ | [1.5-2.1]$\times10^6$ | $[1-5] \times 10^5$ $[1-2] \times 10^5$ | 0 |
| $d^\dagger$ | [0.2541565-0.3558435] | [0.2541565-0.3558435] | [0.2541565-0.3558435] |
| $k_P{}^\dagger$ | [0.0791635-0.1108365] | [0.0791635-0.1108365] | [0.0791635-0.1108365] |
| $x_P{}^\dagger$ | [0.50939629-0.71320371] | [0.50939629-0.71320371] | [0.50939629-0.71320371] |
| $x_N{}^\dagger$ | [0.24932336-0.34907664] | [0.24932336-0.34907664] | [0.24932336-0.34907664] |
| $d_P{}^\dagger$ | [0.049998-0.070002] | [0.049998-0.070002] | [0.049998-0.070002] |
| $d_N{}^\dagger$ | [0.05066464-0.07093536] | [0.05066464-0.07093536] | [0.05066464-0.07093536] |
| $n^\dagger$ | [6.6664-9.3336] | [6.6664-9.3336] | [6.6664-9.3336] |
| $k_I$ | [5.3-6.93]$\times10^8$ | [0.863-2]$\times10^9$ | [1.18-1.21]$\times10^9$ |
| $\gamma$ | [0.99996-1.40004] | [0.74-0.8] | [1.65-1.8] |
| $g_0$ | [0.03-0.06] | [0.208325-0.291675] | [0.26-0.3] |
| $g_0$ | [0.0125-0.0175] | [0.0125-0.0175] | [0.0125-0.0175] |
| $T_{N0}$ | [0.8333-1.1667] | [0.8333-1.1667] | [0.8333-1.1667] |
| $T_{N0}$ | [0] | [0] | [0] |

$^\dagger$ Constant parameters. Perturbation of ± 16.67%

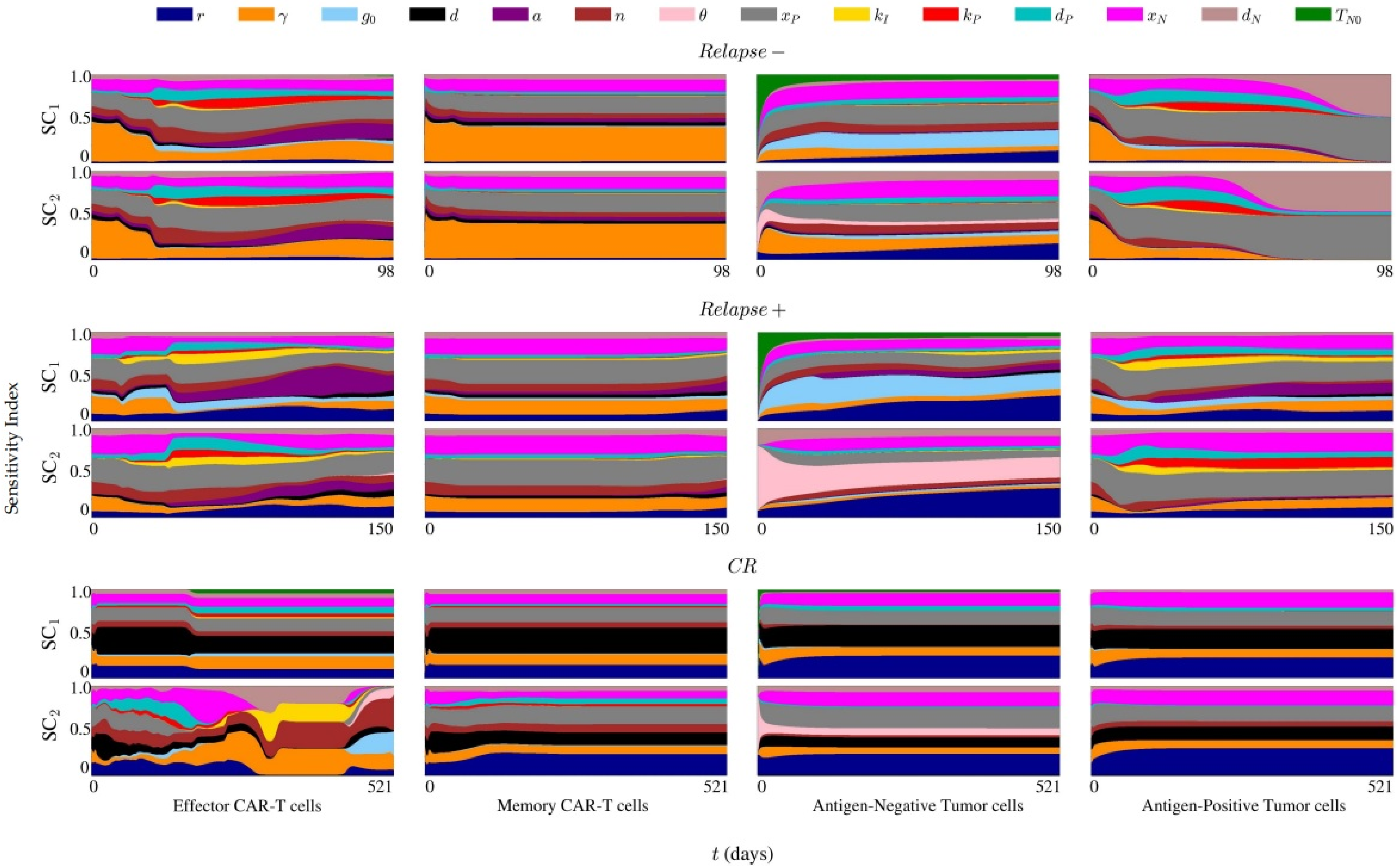


Figure SM-6: Time evolution of the first-order sensitivity index for parameters associated with the tumor dynamics and the resistance mechanisms with respect to each state variable.

#### SM-6 Model Settings and Numerical Simulations

The system of differential equations was uniformly discretized along the antigen expression axis with a step of $\Delta x$= 0.005, and along the temporal axis with a step of $\Delta t = 0.1$ days. The model is solved in the domain $[0,1] \times [(]0, T_{max}]$, in which $T_{max}$ is the maximum simulation time. Given $\Delta x$ and $\Delta t$, the corresponding discretization points are $x_i = i\Delta x, i = 0, \ldots, N_x$ and $t_j = j\Delta t, j = 0, \ldots, N_t$, with $N_x = 1/\Delta x$ and $N_t = T_{max}/\Delta t$. The set of equations is solved iteratively in a decoupled way for each time step. We denote by $k$ the iteration index. The partial differential of antigen-positive tumor cells was solved by combining a central finite difference scheme for the antigen-expression derivatives and the Newmark method to integrate over time. The ordinary differential equations were then solved via a classic fourth-order Runge-Kutta method until convergence was attained. In appropriate units, the initial conditions for tumor cells are

$$T_P(x,0) = \frac{T_{P0}}{d_P\sqrt{2\pi}} e^{-\frac{1}{2}\left(\frac{x-x_P}{d_P}\right)^2} \quad \text{and} \quad T_N(x,0) = \frac{T_{N0}}{d_N\sqrt{2\pi}} e^{-\frac{1}{2}\left(\frac{x-x_N}{d_N}\right)^2} \;, \tag{3}$$

and for CAR-T cells

$$C_T(0) = \text{patient-specific value} \quad \text{and} \quad C_M(0) = 0 \;, \tag{4}$$

where $T_{P0}$ and $T_{N0}$ are the initial number of antigen-positive and antigen-negative tumor cells, respectively. The initial conditions are displayed in Table SM-4.

The boundary conditions at $x = 0$ and $x = 1$ impose zero cell flux and are given by:

$$v(x,t)T_P(x,t) - \sigma \frac{\partial T_P(x,t)}{\partial x} = 0 \quad \text{and} \quad \frac{\partial T_N(x,t)}{\partial x} = 0 \;. \tag{5}$$

##### SM-6.1 Numerical Methods

For a generic state variable $\bar{T}(x,t)$, we denote

$$\bar{T}^t(x) = \bar{T}^t(x_i) = \bar{T}^t_i, \quad \bar{T}^t(x-\Delta x) = \bar{T}^t(x_{i-1}) = \bar{T}^t_{i-1}, \quad \text{and} \quad \bar{T}^t(x+\Delta x) = \bar{T}^t(x_{i+1}) = \bar{T}^t_{i+1}. \tag{6}$$

Using the second-order centered finite difference method, we can define the following approximations for the space derivatives:

$$\frac{\partial \bar{T}^t(x)}{\partial x} = \frac{\bar{T}^t_{i-1} - \bar{T}^t_{i+1}}{2\Delta x}, \quad \text{and} \quad \frac{\partial^2 \bar{T}^t(x)}{\partial^2 x} = \frac{\bar{T}^t_{i-1} - 2\bar{T}^t_i + \bar{T}^t_{i+1}}{\Delta x^2} \;. \tag{7}$$

Now, using the notation $\frac{\partial \bar{T}(x,t)}{\partial t} = \dot{\bar{T}}^t_i$, $\frac{\partial^2 \bar{T}(x,t)}{\partial^2 t} = \ddot{\bar{T}}^t_i$, we apply the Newmark method [35] to integrate over time and get:

$$\dot{\bar{T}}^t_i = a_0 a_7 (\bar{T}^t_i - \bar{T}^{t-\Delta t}_i) + (1 - a_7 a_2)\dot{\bar{T}}^{t-\Delta t}_i + (a_6 - a_7 a_3)\ddot{\bar{T}}^{t-\Delta t}_i \tag{8}$$

with:

$$a_0 = \frac{1}{\beta \Delta t^2},\ a_1 = \frac{\gamma}{\beta \Delta t},\ a_2 = \frac{1}{\beta \Delta t},\ a_3 = \frac{1}{2\beta} - 1,\ a_4 = \frac{\gamma}{\beta} - 1,\ a_5 = \frac{\Delta t}{2}\left[\frac{\gamma}{\beta} - 2\right],\ a_6 = \Delta t(1-\gamma),\ a_7 = \gamma \Delta t, \tag{9}$$

where $\gamma$ and $\beta$ are parameters of the method, selected according to stability, precision, and efficiency criteria. There are many variations of this algorithm, such as the linear acceleration method scheme ($\gamma$=1/2 and $\beta$=1/6). Here we use the original Newmark method, which is an unconditionally stable scheme known as the trapezoidal rule, where $\gamma$=1/2 and $\beta$=1/4.

To approximate the ordinary differential equations that model CAR-T cell populations, we introduce a new state variable $\bar{C}(t)$ and define the general equation

$$\frac{\partial \bar{C}(t)}{\partial t} = f(t, \bar{C}) \;, \tag{10}$$

where $f(t, \bar{C})$ denotes a function of $\bar{C}$ at time $t$. Using the classic fourth-order Runge-Kutta method, we have

$$\bar{C}^t = \bar{C}^{t-\Delta t} + \frac{1}{6}(k_1 + 2k_2 + 2k_3 + k_4) \; , \tag{11}$$

where

$$\begin{aligned} k_1 &= f(t-\Delta t, \bar{C}^{t-\Delta t}), \quad k_2 = f\left(t - 0.5\Delta t, \bar{C}^{t-\Delta t} + 0.5\Delta t k_1\right) \; , \\ k_3 &= f\left(t - 0.5\Delta t, \bar{C}^{t-\Delta t} + 0.5\Delta t k_2\right) , \quad k_4 = f(t, \bar{C}^{t-\Delta t} + \Delta t k_3) \; . \end{aligned} \tag{12}$$

###### SM-6.1.1 Antigen-positive tumor population

Applying the chain rule on the advective term of Equation (2), we may rewrite it in the form

$$\frac{\partial T_P(x,t)}{\partial t} = -v(x,t)\frac{\partial T_P(x,t)}{\partial x} + \sigma\frac{\partial^2 T_P(x,t)}{\partial x^2} + T_P(x,t)\left[-k_P + r(1-\theta) - \gamma g(x) f(C_T(t), T(t))\right] \; . \tag{13}$$

Now, applying the previously defined approximations, the semi-discrete form of Equation 13 in the $k^{th}$ iteration is given by

$$\underbrace{\frac{\partial T_P(x,t)}{\partial t}}_{\text{Newmark}} = -\underbrace{v(x,t)}_{\text{at } k-1}\left[\frac{T^t_{P_{i-1}} - T^t_{P_{i+1}}}{2\Delta x}\right] + \sigma\left[\frac{T^t_{P_{i-1}} - 2T^t_{P_i} + T^t_{P_{i+1}}}{\Delta x^2}\right] + T^t_i\left[-k_P + r(1-\theta) - \gamma g(x)\underbrace{f(C_T(t), T(t))}_{\text{at } k-1}\right] ,$$

or, similarly,

$$\underbrace{\frac{\partial T_P(x,t)}{\partial t}}_{\text{Newmark}} = v^{k-1}(x_i,t)\left[\frac{T^t_{P_{i+1}} - T^t_{P_{i-1}}}{2\Delta x}\right] + \sigma\left[\frac{T^t_{P_{i-1}} - 2T^t_{P_i} + T^t_{P_{i+1}}}{\Delta x^2}\right] + T^t_i\left[-k_P + r(1-\theta) - \gamma g_i f^{k-1}(C^t_T, T^t)\right] ,$$

in which we drop the index of the current iteration to ease notation. Applying Equation 8 of the Newmark method and separating the terms in $T_P$ evaluated at the previous time $t - \Delta t$ from those at the current time $t$, we obtain:

$$\begin{aligned} &\underbrace{\left[\frac{2}{\Delta t}\right](T^{t-\Delta t}_{P_i}) - \dot{T}^{t-\Delta t}_{P_i}}_{\mathbf{b}} = \\ &\underbrace{v^{k-1}(x_i,t)\left[\frac{T^t_{P_{i+1}} - T^t_{P_{i-1}}}{2\Delta x}\right] + \sigma\left[\frac{T^t_{P_{i-1}} - 2T^t_{P_i} + T^t_{P_{i+1}}}{\Delta x^2}\right] + T^t_{P_i}\left[-k_P + r(1-\theta) - \gamma g_i f^{k-1}(C^t_T, T^t) - \left[\frac{2}{\Delta t}\right]\right]}_{\mathbf{AT_P}} . \end{aligned} \tag{14}$$

Assembling the contribution of each discretization point in the antigen expression domain we build the system of algebraic equations $\mathbf{AT_P} = \mathbf{b}$ which is solved by using the LU decomposition method.

###### SM-6.1.2 Antigen-negative tumor population

We now apply the previously defined approximations into Equation (3) that models the dynamics of $T_N(x,t)$ to get

$$\underbrace{\frac{\partial T_N(x,t)}{\partial t}}_{\text{Newmark}} = rT^t_{N_i} + \theta r W_i \int_0^1 \underbrace{T_P(x,t)}_{\text{at } k-1} dx - T^t_{N_i}\left[\gamma g(x)\underbrace{f(C_T(t), T(t))}_{\text{at } k-1}\right] .$$

Using Equation 8 of the Newmark method yields

$$\underbrace{\left[\frac{\gamma}{\beta\Delta t}\right]}_{2/\Delta t}(T^t_{N_i} - T^{t-\Delta t}_{N_i}) + \underbrace{\left[1 - \frac{\gamma}{\beta}\right]}_{-1}\dot{T}^{t-\Delta t}_{N_i} + \underbrace{\left[\Delta t(1-\gamma) - (\gamma\Delta t)\left[\frac{1}{2\beta} - 1\right]\right]}_{0}\ddot{T}^{t-\Delta t}_{N_i} =$$

$$rT^t_{N_i} + \underbrace{\theta r W_i \int_0^1 T^{k-1}_{P_i} dx}_{\text{Constant } = \mathcal{C}^{k-1}} - T^t_{N_i}\left[\gamma g_i f^{k-1}(C^t_T, T^t)\right] .$$

Rearranging the previous equation, we finally obtain the antigen-negative tumor population by solving

$$T^t_{N_i} = \frac{-\left[\frac{2}{\Delta t}\right]T^{t-\Delta t}_{N_i} + \dot{T}^{t-\Delta t}_{N_i} - \mathcal{C}^{k-1}}{\left[r - \left[\frac{2}{\Delta t}\right] - \gamma g_i f^{k-1}(C^t_T, T^t)\right]} . \tag{15}$$

##### SM-6.1.3 CAR-T cell populations

The effector and memory CAR-T cells, respectively given by Equations (11) and (12), can be rewritten as

$$\frac{\partial C_T(t)}{\partial t} = \underbrace{\kappa^t\left[\frac{T^t_S}{A + T^t_S}\right]C^t_T - \mu C^t_T - \epsilon C^t_T + \theta_M C^t_M T^t_S - \alpha C^t_T T(t)}_{f(t, C_T)} , \tag{16}$$

$$\frac{\partial C_M(t)}{\partial t} = \underbrace{\epsilon C^t_T - \theta_M C^t_M T^t_S - \mu_M C^t_M}_{f(t, C_M)} , \tag{17}$$

and, through the application of Equations 11 and 12, we find $C_T(t)$ and $C_M(t)$.

#### SM-6.2 Convergence criteria

The convergence of the nonlinear process is attained only when all state variables satisfy:

$$\begin{aligned} \delta_{T_P} &= \frac{\|T^{k+1}_{Pi} - T^k_{Pi}\|}{\|T^k_{Pi}\|} \leq tol \ , \ \delta_{T_N} = \frac{\|T^{k+1}_{Ni} - T^k_{Ni}\|}{\|T^k_{Ni}\|} \leq tol \ , \\ \delta_{C_T} &= \frac{\|C^{k+1}_T - C^k_T\|}{\|C^k_T\|} \leq tol \ , \text{ and } \delta_{C_M} = \frac{\|C^{k+1}_M - C^k_M\|}{\|C^k_M\|} \leq tol \ , \end{aligned} \tag{18}$$

where $tol = 10^{-8}$ is the adopted tolerance.

#### SM-6.3 Solution algorithm

The proposed system of integro-differential equations that describes the interplay among antigen-positive tumor cells, antigen-negative tumor cells, effector CAR-T cells, and memory CAR-T cells is solved through the steps defined in the following algorithm:

**Algorithm** Solution to the integro-partial differential model

Set model parameters, numerical constants, and initial and boundary conditions
$j \leftarrow 1$
**while** $j\Delta t \leq T_{max}$ **do**
  Update the variables with the solution of the previous time step
  **while** $\delta_{T_P}, \delta_{T_N}, \delta_{C_T}, \delta_{C_M} > tol$ **do**
    $i \leftarrow 0$
    **while** $i\Delta x \leq 1$ **do**
      Solve Equation 15 for $T_N(x_i, t_j)$
      Assemble in $\mathbf{A}$ and $\mathbf{b}$ the contribution from Equation 14 for $T_P(x_i, t_j)$
      $i \leftarrow i + 1$
    **end while**
    Solve the system of equations $\mathbf{AT_P} = \mathbf{b}$ for $\mathbf{T_P}(x, t_j)$
    Solve Equations 16 and 17 for $C_T(t_j)$ and $C_M(t_j)$
  **end while**
  Update the variables with the solution of the last iteration
  $j \leftarrow j + 1$
**end while**
